# PTH regulates osteogenesis and suppresses adipogenesis through Zfp 467 in a feed-forward, cyclic AMP-dependent manner

**DOI:** 10.1101/2022.10.10.511580

**Authors:** Hanghang Liu, Akane Wada, Isabella Le, Phuong T Le, Andrew WF Lee, Jun Zhou, Francesca Gori, Roland Baron, Clifford J. Rosen

## Abstract

Conditional deletion of the PTH1R in mesenchymal progenitors reduces osteoblast differentiation, enhances marrow adipogenesis and increases zinc finger protein 467 (*Zfp467*) expression. In contrast, genetic loss of *Zfp467* increased *Pth1r* expression and shifts mesenchymal progenitor cell fate towards osteogenesis and higher bone mass. In this study we hypothesized that PTH1R and ZFP467 could constitute a feedback loop that facilitates PTH-induced osteogenesis and that conditional deletion of *Zfp467* in osteogenic precursors would lead to high bone mass. We report that *PrrxCre Zfp467* but not *AdipoCre Zfp467* mice exhibit an identical phenotype to the *Zfp467*-/-mice with high bone mass and greater osteogenic differentiation We also found that PTH suppressed *Zfp467* expression primarily via the cyclic AMP/PKA pathway. Not surprisingly, PKA activation inhibited the expression of *Zfp467* and gene silencing of *Pth1r* caused an increase in *Zfp467* mRNA transcription. Dual fluorescence reporter assays and confocal immunofluorescence demonstrated that genetic deletion of *Zfp467* resulted in higher nuclear translocation of p50 that binds to the P2 promoter of the *Pth1r* and increased its transcription. As expected, *Zfp467-/-* cells had enhanced production of cyclic AMP and increased glycolysis in response to exogenous PTH. Additionally, the osteogenic response to PTH was also enhanced in *Zfp467* -/-calvarial osteoblasts, and the pro-osteogenic effect of *Zfp467* deletion was blocked by gene silencing of *Pth1r* or a PKA inhibitor. In conclusion, our findings suggest that loss or PTH1R-mediated repression of *Zfp467* results in a pathway that increases *Pth1r* transcription via p50 and thus cellular responsiveness to PTH, ultimately leading to enhanced bone formation.

## Introduction

Intermittent administration of parathyroid hormone (PTH 1-34), teriparatide, is anabolic for bone and is a well-established treatment for osteoporosis (1). Both PTH and PTHrp act through the PTH1R to drive bone formation, increase bone mass and lower fracture risk (2). It has been established that one mechanism for PTH-driven osteogenesis is through induction of cyclic AMP and the PKA pathway, although PKC can also be activated by ligand binding to the PTH1R (3). Downstream targets from PTH1R activation include IGF-1, FGF-2, Rankl, Sclerostin, Wnt signaling, salt-inducible kinases (SIKs) and the BMPs(3-5). *Pth1R* is expressed in chondrocytes, and the entire osteoblast lineage from early osteogenic progenitors to osteoblasts during which it is up-regulated (2). Osteocytes and lining cells also express *pth1r* (2,3). Recent studies have noted that the PTH1R is also expressed in mature adipocytes and their immediate precursors, marrow adipocyte like progenitors or MALPs (6).

Intermittent PTH treatment increases bone formation both by enhancing the number of osteoblasts and their function, resulting in higher bone mass (7). Several reports have demonstrated that PTH-induced lineage allocation of mesenchymal skeletal stromal cells into osteoblasts is at the expense of adipogenesis (8). This is consistent with human studies which confirmed that PTH treatment reduces bone marrow adiposity principally through a shift in lineage allocation (9). The transcriptional mechanisms whereby PTH drives a progenitor cell towards an osteoblast are multiple, complex and redundant. Previously we reported that genetic deletion of the PTH1R in Prx1^+^ cells, resulted in low bone mass, and a marked increase in bone marrow adiposity(8). In our search for the mechanisms that drive adipogenesis we identified *Zfp467* as a gene markedly upregulated in the absence of the PTH1R(8). Zinc finger proteins (ZFPs) are one of the largest classes of transcription factors in eukaryotic genomes and *Zfp521* and *Zfp423* have been identified as critical determinants of adipogenesis (10,11). *Zfp467* has been reported to enhance both *Sost* and *Pparg* expression in marrow stromal cells, and is highly expressed in both adipocyte and osteoblast progenitors(12). And, an earlier study identified *Zfp467* as a transcriptional regulator of lineage allocation among mesenchymal cells of the marrow(13). In accordance with that report, we reported that global genetic deletion of *Zfp467* resulted in high bone mass and a reduction in bone marrow adipose tissue and peripheral adipose depots(14). Hence the inverse relationship between PTH1R and *Zfp467* suggested a pathway by which PTH could impact lineage allocation. To determine the molecular mechanism of this interaction, we studied *Zfp467*-/-osteoblast progenitors and identified a potential transcriptional modulator of the PTH1R, regulated by ZFP467. This report details both in vitro and in vivo studies, and points to a novel pathway that impacts skeletal formation through the PTH1R.

## Results

Previously, we showed that global deletion of *Zfp467* increased trabecular bone volume and cortical bone thickness, compared to wild-type mice at the age of 16 weeks (14). And histomorphometry results showed higher structural and dynamic formation parameters in *-/-* mice vs. +/+(14). To assess whether the effect on bone mass seen in the *Zfp467* global KO mice was cell-autonomous to mesenchymal cells, we generated mice with deletion of Zfp467 in the limb MSCs by crossing *Zfp467*^fl/fl^ mice with the *Prrx1Cre* mice and *AdiponectinCre* mice. (**Fig.1-figure supplement 1**)

### *Prrx Cre Zfp467* mice have reduced cortical and trabecular bone mass and recapitulate the global Zfp467 null mice

Body mass and body size were not significantly different between *Prrx1Cre; Zfp467*^*fl/fl*^ and control littermates in both males and females (data not shown). μCT analysis showed that *Prrx1Cre; Zfp467*^*fl/fl*^ mice had higher trabecular bone volume fraction (Tb.BV/TV), greater connectivity density (Conn.D), and higher trabecular number (Tb.N) with a significant decrease in structural model index (SMI) and trabecular separation (Tb.Sp) in both males and females (Fig.1 A, B), indicating an increase in trabecular bone mass. Cortical thickness was increased, although not significantly in males and decreased in females. In contrast, tibial adipose tissue volume fraction in the marrow (Ad.V/TV) was significantly decreased in males (Fig. 1A), and showed a similar non-significant trend in females (Fig. 1B). Although the time point of examination differed (12 week instead of 16 week), overall, these data established that skeletal changes observed in the global *Zfp467* KO mice were mostly attributable to changes in the MSC lineage.

**Fig. 1.**
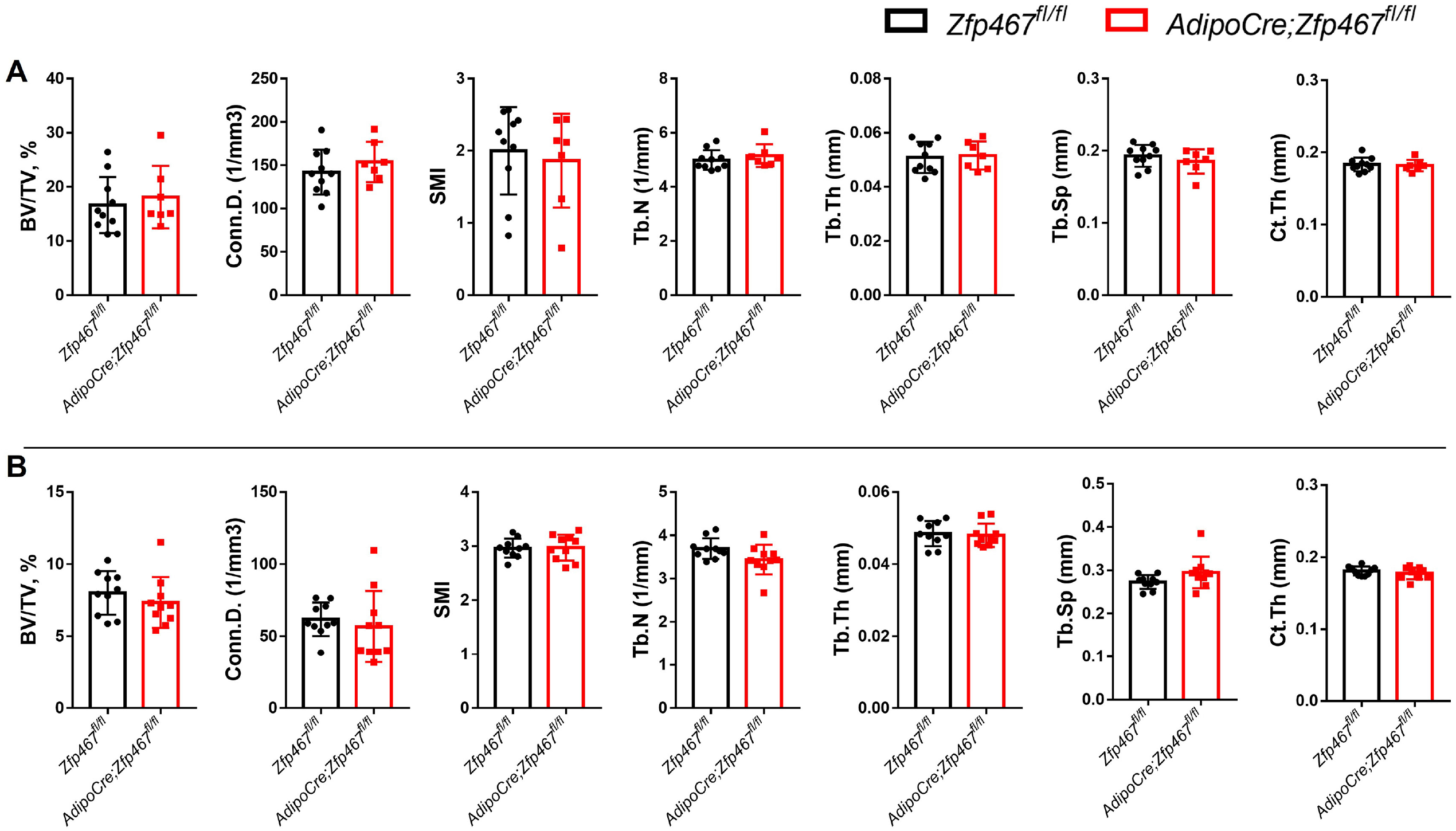
*Prrx1 Cre Zfp467* mice have more trabecular bone mass and less adipose tissue in bone marrow, recapitulating the global Zfp467 null mice: (A) Male and (B) Female 12 weeks old *Prrx1Cre; Zfp467fl/fl* mice and control mice were measured representative μCt images of tibia trabecular and cortical bone of tibiae. Marrow adipose tissue volume (Ad.V) was quantified by osmium tetroxide staining and μCT. Data shown as mean ± SD by unpaired Student’s t test, n=5-8 per group.

### *Adiponectin Cre Zfp467* mice have similar cortical and trabecular bone mass to controls

We initially sought to determine the whole body phenotype of the conditional knock out of *Zfp467* mice. We found that *Zfp467*^*fl/fl*^; *AdipoCre* mice have similar body weight, fat mass, lean mass and femoral areal BMD at 12 weeks compared to control mice in both males and females (data not shown). μCT was performed and analyzed in the metaphysis and cortical bone at the tibial mid-diaphysis in 12-week-old *Zfp467*^*fl/fl*^; *AdipoCre* mice and control mice. Not surprisingly, no significant difference was found regarding trabecular bone volume fraction (BV/TV), trabecular bone mineral density (BMD), connectivity density (Conn.D), trabecular number (Tb.N), trabecular thickness (Tb.Th), trabecular separation (Tb.Sp) and structural model index (SMI) (**Fig .1-figure supplement 2** A, B) between controls and AdipoCre mice. In addition, we did not observe any differences in cortical bone measurements including total area (Tt. Ar), cortical area/total area (Ct.Ar/Tt.Ar), cortical thickness (Ct.Th), marrow area (Ma.Ar), cortical porosity (Ct. Porosity) and cortical tissue mineral density (Ct.TMD) (**Fig .1-figure supplement 2** A, B).

### PTH suppressed the expression levels of *Zfp467* via the PKA pathway

To understand the mechanisms that underly the impact of mesenchymal deletion of Zfp467, we first examined the signaling pathway of PTH relative to Zfp467 in osteoblasts. Consistent with a previous study which showed that short term PTH treatment suppressed the expression of *Zfp467*(12), our study confirmed that treating cells with 100 nM PTH for 10min could significantly suppress *Zfp467* expression in both COBs (**Fig. 2A**) and BMSCs (**Fig. 2B**). When pretreating COBs and BMSCs with PKA and PKC inhibitors (10uM H89 and 5uM Go6983, Selleck Chemicals, Houston, TX), respectively, for 2 hours prior to 10min of 100 nM PTH treatment, significant rescue of *Zfp467* suppression was seen in H89 group, but Go6983 had no effect (**Fig. 2A, B**). Forskolin, a selective PKA pathway activator was also found to significantly inhibit the expression level of *Zfp467* in COBs (**Fig. 2C**) and BMSCs (**Fig. 2D**). Moreover, consistent with our previous study that *Prx1*-*Cre Pth1r*^*fl/fl*^ mice showed higher expression level of *Zfp467* in bone marrow, we silenced *Pth1r* with siRNA in both COBs and BMSC and found *Pth1r* knock-down significantly up-regulated the expression level of *Zfp467* (**Fig. 2E, F**). These data suggested that PTH1R activation could down-regulate *Zfp467* expression via PKA pathway, and that ZFP467 could be one of the important downstream targets of PTH signaling.

**Fig. 2.**
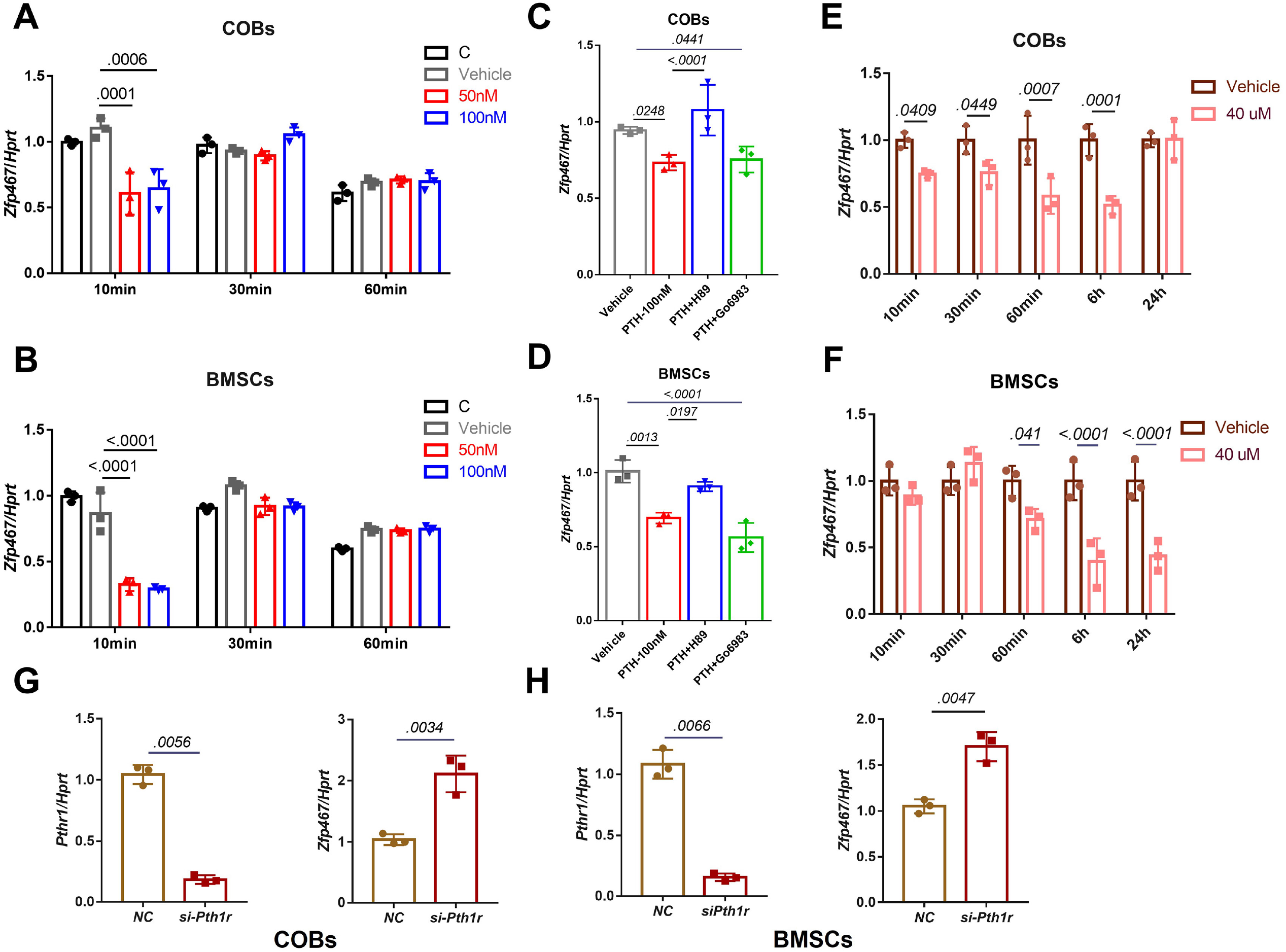
PTH suppressed the expression levels of *Zfp467* via the PKA pathway. (A) PTH treatments significantly suppressed *Zfp467* expression within 10 minutes of treatment in *+/+* COBs. Data shown as mean ± SD by one-way ANOVA, n=3 independent experiments for each group. (B) PTH treatments significantly suppressed *Zfp467* expression within 10 minutes of treatment in *+/+* BMSCs. Data shown as mean ± SD by one-way ANOVA, n=3 independent experiments for each group. (C) qPCR results· of *Zfp467+/+* in COBs with 2 hours PKA or PKC inhibitor treatment prior to 10min of 100 nM PTH exposure, PKA but not PKC inhibitor was able to rescue the suppression of *Zfp467* induced by PTH. Data shown as mean ± SD by one-way ANOVA, n=3 independent experiments for each group. (D) qPCR results of *Zfp467+/+* in BMSCs with 2 hours PKA or PKC inhibitor treatment prior to 10min of 100 nM PTH exposure. PKA but not PKC inhibitor was able to rescue the suppression of *Zfp467* induced by PTH. Data shown as mean ± SD by one-way ANOVA, n=3 independent experiments for each group. (E) Forskolin significantly suppressed *Zfp467* expression within 1-hour of treatment in +/+ COBs. Data shown as mean ± SD by unpaired Student’s t test, n=3 independent experiments for each group. (F) Forskolin significantly suppressed *Zfp467* expression after 6-hours of treatment in +/+ BMSCs. Data shown as mean ± SD by unpaired Student’s t test, n=3 independent experiments for each group. (G) *Pth1r-*siRNA treatment in +/+ COBs led to an increase of *Zfp467* expression in +/+ COBs. Data shown as mean ± SD by unpaired Student’s t test, n=3 independent experiments for each group. (H) *Pth1r-*siRNA treatment in +/+ COBs led to an increase of *Zfp467* expression in +/+ BMSCs. Data shown as mean ± SD by unpaired Student’s t test, n=3 independent experiments for each group.

### *Zfp467-/-* cells have greater *Pth1r* transcriptional levels driven by both the P1 and P2 promoter

Our previous study showed that the global absence of *Zfp467* resulted in a significant increase in trabecular bone volume, a marked reduction in peripheral and marrow adipose tissue, and a ∼40% increase in *Pth1r* expression in bone from the *-/-* mice compared to littermate controls(14). These data suggested the possibility of a positive feedback loop whereby the suppression of *Zfp467* mediated by PTH leads to an increase of PTH1R. Consistent with this tenet, we found higher gene and protein expression level of PTH1R in *Zfp467-/-* COBs and BMSCs (**Fig.3A, B**). Three transcripts of *Pth1r* (NM_011199.2, NM_001083935.1 and NM_001083936.1) with different transcription starting sites (TSS) were reported based on NCBI database and UCSC Genome Browser on Mouse (**Fig. 3C**). The three transcripts shared the same coding sequence and the only difference was located at the three prime untranslated region. Based on the different 3 prime untranslated regions, we designed related primers; by qPCR results both the *Pth1r-T1* and *Pth1r-T2* transcripts were upregulated in *Zfp467-/-* cells (**Fig. 3D**). *Pth1r-T1* and *Pth1r-T2* transcripts were driven by P1 and P2 promoters, respectively. As P1 is much longer that P2 and hadn’t been investigated before, we designed a P1 promoter-driven dual-fluorescence reporter with four different length P1 promoter to assess any change in the promoter activity of P1 in both COBs and BMSCs in the absence of 467 (**Fig. 3C**). We found that the 1.6 kb and 2.1 kb promoter-driven reporters are higher in activated -/- cells compared to +/+ cells (**Fig. 3E**), which indicated the binding site of *Pth1r* P1 promoter in -/- cells is between 0.6 kb and 1.1 kb ahead of P1 TSS. Combined with the previously reported two potential transcription factor binding sites of P2(15), we constructed three P1 and P2 driven dual-fluorescence reporters (**Fig. 3F**). Both P1 and P2 were activated in -/- cells, but P2 showed much higher activity in BMSCs and COBs. P2-2 was also much more active in -/- cells than +/+ cells. P2-2 was therefore chosen for transcription factor prediction via JASPAR, AnimalTFDB and PROMO database and six transcription factors were noted, including CREB, EBF1, MYOD, cFOS, p50 and GATA1.

**Fig. 3.**
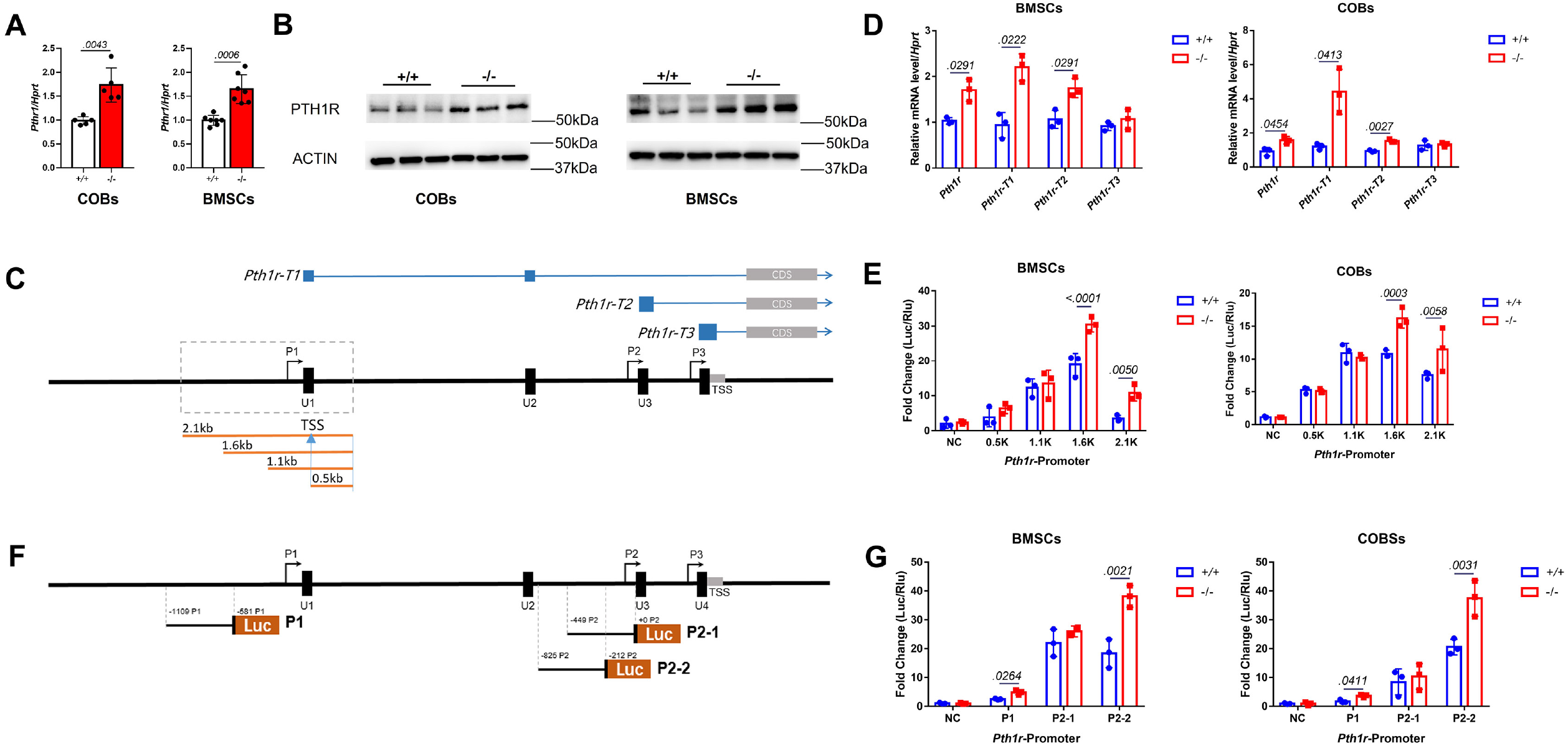
*Zfp467-/-* cells have greater *Pth1r* transcriptional levels driven by both the P1 and P2 promoter. (A) qPCR results of baseline COBs and BMSCs. Higher expression level of *Pth1r* was found in both *-/-* COBs and BMSCs; Data shown as mean ± SD by unpaired Student’s t test, n=5-7 independent experiments for each group. (B) Western blot analysis of baseline COBs and BMSCs. Higher expression level of PTH1R was found in both *-/-* COBs and BMSCs; (C) A schematic of three different *Pth1r* transcripts and P1 promoter of *Pth1r*. Four different length P1 promoter constructs were designed and inserted into dual-fluorescence reporter vector. (D) qPCR results of three *Pth1r* transcripts and total *Pth1r*. Total *Pth1r* and *Pth1r-T1, T2* but not *Pth1r-T3* were up-regulated in both -/- COBs and BMSCs. Data shown as mean ± SD by unpaired Student’s t test, n=3 independent experiments for each group. (E) Dual-fluorescence assay using indicated four P1 reporter constructs. 1.6 kb and 2.1 kb constructs driven reporter is higher activated in -/- cells compared to +/+ cells. Data shown as mean ± SD by unpaired Student’s t test, n=3 independent experiments for each group. (F) A schematic of P1 and P2 promoter constructs of *Pth1r*. (G) Dual-fluorescence assay using indicated P1 and P2 reporter constructs. Both P1 and P2-2 were found significantly higher activated in *-/-* cells. Data shown as mean ± SD by unpaired Student’s t test, n=3 independent experiments for each group.

### *Zfp467-/-* cells have higher p50 and GATA1 nuclear translocation

Using qPCR, we found no differences in the transcriptional levels of these transcription factors (**Fig.4-figure supplement**). Therefore, we overexpressed all of the potential transcription factors that may bind to Pth1r P2-2 promoter in a pre-osteoblast cell line MC3T3-E1, and only GATA1 and p50 overexpression could significantly upregulate the expression level of *Pth1r*, especially *Pth1r-T1* and -*T2*(**Fig. 4A**). We then detected the nuclear translocation level of p50 and GATA1 using immune-florescence, nuclear protein isolation and western blot. p50 and GATA1 were almost evenly distributed in the cytoplasm and nucleus of +/+ cells but underwent partial translocation to the nucleus in *Zfp467 -/-* cells (**Fig. 4B**). *Zfp467* -/- cells also showed much higher nuclear protein level of both p50 and GATA1 (**Fig. 4C, D**).

**Fig. 4.**
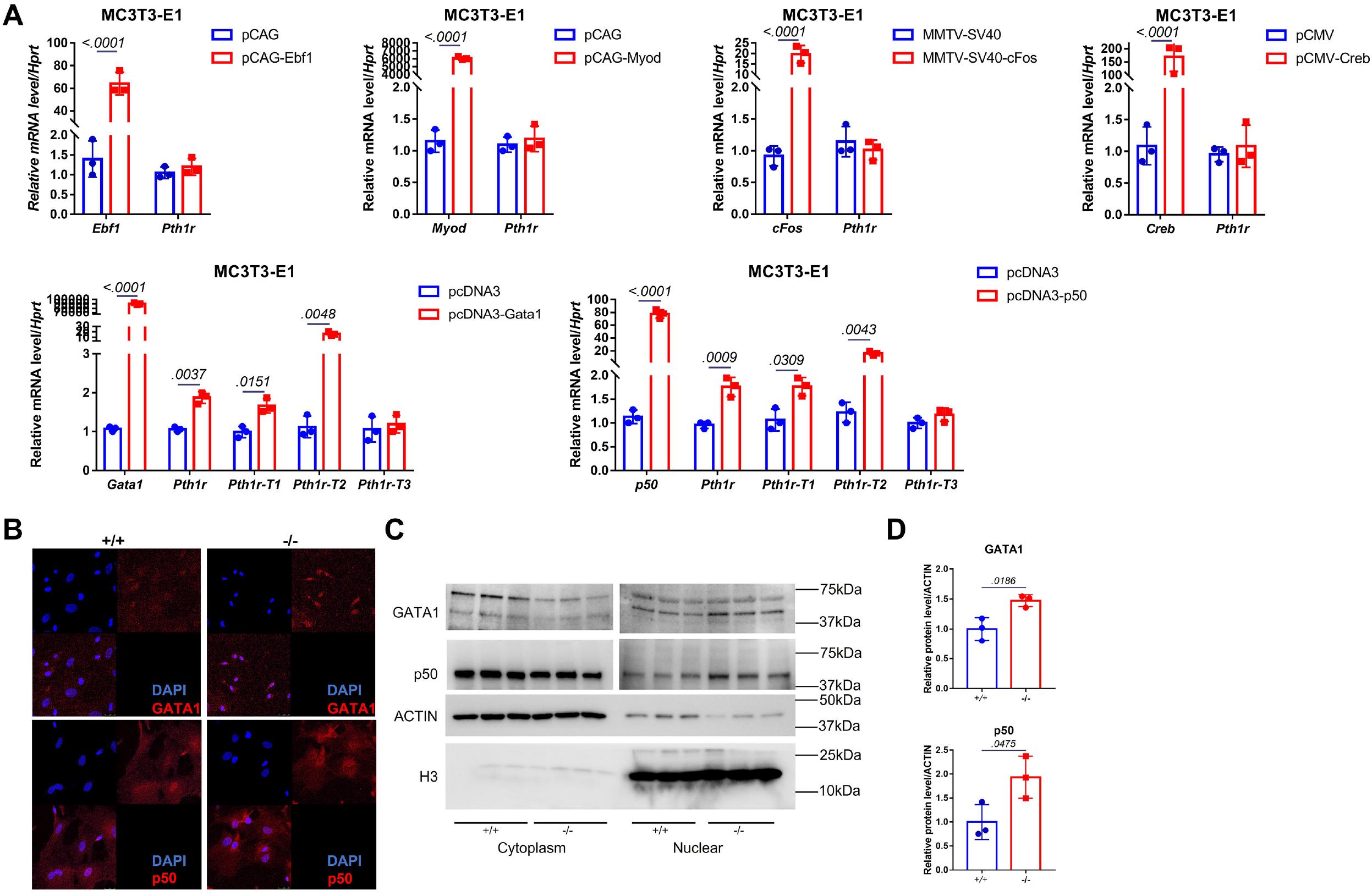
*Zfp467-/-* cells have higher p50 and GATA1 nuclear translocation. (A) qPCR results of overexpression of *Ebf1, Myod, Myog, Gata1* and *p50* in MC3T3-E1 cell line. GATA1 and p50 overexpression could significantly upregulate the expression level of *Pth1r*. Data shown as mean ± SD by unpaired Student’s t test, n=3 independent experiments for each group. (B) Representative confocal images of GATA1 and p50 immune-florescence in *Zfp467+/+* and *Zfp467-/-* BMSCs. (C) Nuclear protein level of GATA1 and p50 in *Zfp467+/+* and *Zfp467-/-* BMSCs. (D) Quantification analysis for nuclear protein level of GATA1 and p50 in *Zfp467+/+* and *Zfp467-/-* BMSCs. Data shown as mean ± SD, n=3 independent experiments for each group. Data shown as mean ± SD by unpaired Student’s t test, n=3 independent experiments for each group.

### p50-Relb heterodimer may drive greater *Pth1r* transcription in *Zfp467-/-* cells

In order to confirm whether p50 or GATA1 could activate the specific P1 or P2-2 Pth1r promoter, we co-transfected MC3T3-E1 cells with P1 and P2-2 dual-luciferase reporter and GATA1, p50 overexpression plasmids. Only the p50 overexpression group was able to significantly activate the P2-2 promoter (**Fig. 5A**). ChIP results showed that the DNA was properly sheared and IP was successfully conducted (**Fig. 5B**). ChIP-qPCR results showed that the first two parts of P2 were properly enriched in our IP product (>0.5%) (**Fig.5-figure supplement A**), and the first part of P2 was approximately 20 fold more highly enriched in our p50 IP product than IgG (**Fig. 5C, Fig.5-figure supplement B**); this indicated that p50 binds to the P2 promoter, especially at the first 200 bp site. Subsequently, we treated COBs and BMSCs with *p50* siRNA and found that *p50* knock-down could significantly inhibit the expression of *Pth1r* in both *+/+* and *-/-* COBs and BMSCs. Importantly, *p50* knock-down in *-/-* cells reverts the levels of Pth1r to the levels seen in *+/+* cells (**Fig. 5D-F**).

**Fig. 5.**
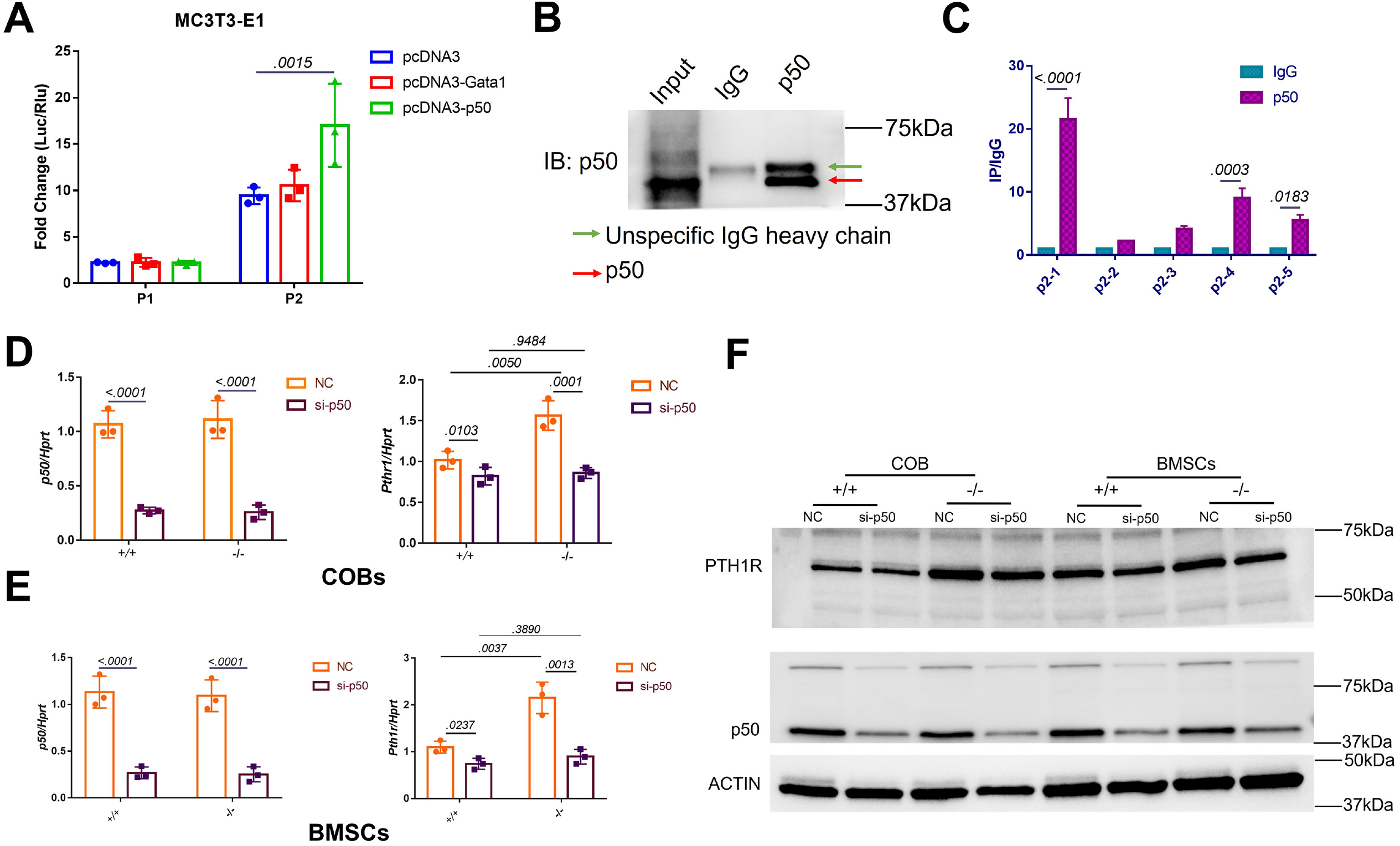
p50 was found to transactive *Pth1r* expression in *Zfp467 -/-* cells. (A) Reporter assays using the indicated P1 or P2 reporter construct and an expression vector bearing Gata1, p50 or a control empty vector. Data shown as mean ± SD by one-way ANOVA, n=3 independent experiments for each group. Data shown as mean ± SD by one-way ANOVA, n=3 independent experiments for each group. (B) Immune-blot assay using a control rabbit IgG antibody (IgG) or the anti-p50 antibody during chromatin immunoprecipitation assay. (C) DNA enrichment of Pth1r P2 promoter, ratio between p50 and IgG IP products, first part and last two parts of P2 were significantly enriched by p50 antibody. Data shown as mean ± SD by unpaired Student’s t test, n=3 independent experiments for each group. (D, E) qPCR results of the expression levels of *p50* and *Pth1r* in *p50* siRNA-treated *Zfp467 +/+* and *-/-* COBs and BMSCs. Data shown as mean ± SD by two-way ANOVA, n=3 independent experiments for each group. (F) Western blot analysis of *p50* and *Pth1r* in *p50* siRNA-treated *Zfp467 +/+* and *-/-* COBs and BMSCs.

In order to determine whether p50 could bind to the Pth1r P2-2 promoter directly, we performed DNA pulldown assay using biotin-labeled *Pth1r P2* promoter as a probe. As shown in **Fig. 6A**, the biotin-Pth1rP2 group showed a specific band in both MC3T3-E1 nuclear extracts and purified p50 protein, suggesting a direct physical interaction between p50 and Pth1r P2 promoter. However, we noticed that p50 does not have a transcriptional activation domain, so p50 must heterodimerize with other transcription factors in order to increase gene transcription. Using String database and checking published studies, we found 8 candidate that might heterodimerize with p50 to regulate gene transcription: NFYC, NPAS1, Rel, AKAP8, RelA, RelB, ANKRD42 and HDAC1. Using siRNA, we knocked down all these potential p50 partners (**Fig. 6B**), but the upregulated *Pth1r* induced by p50 overexpression could only be dampened by *Npas1* and *Relb* siRNA (**Fig. 6C**). Further co-immunoprecipitation results confirmed that p50 could heterodimerize with RelB only **(Fig. 6D**), which suggested p50-Relb heterodimer may drive greater *Pth1r* transcription in Zfp467-/-cells.

**Fig. 6.**
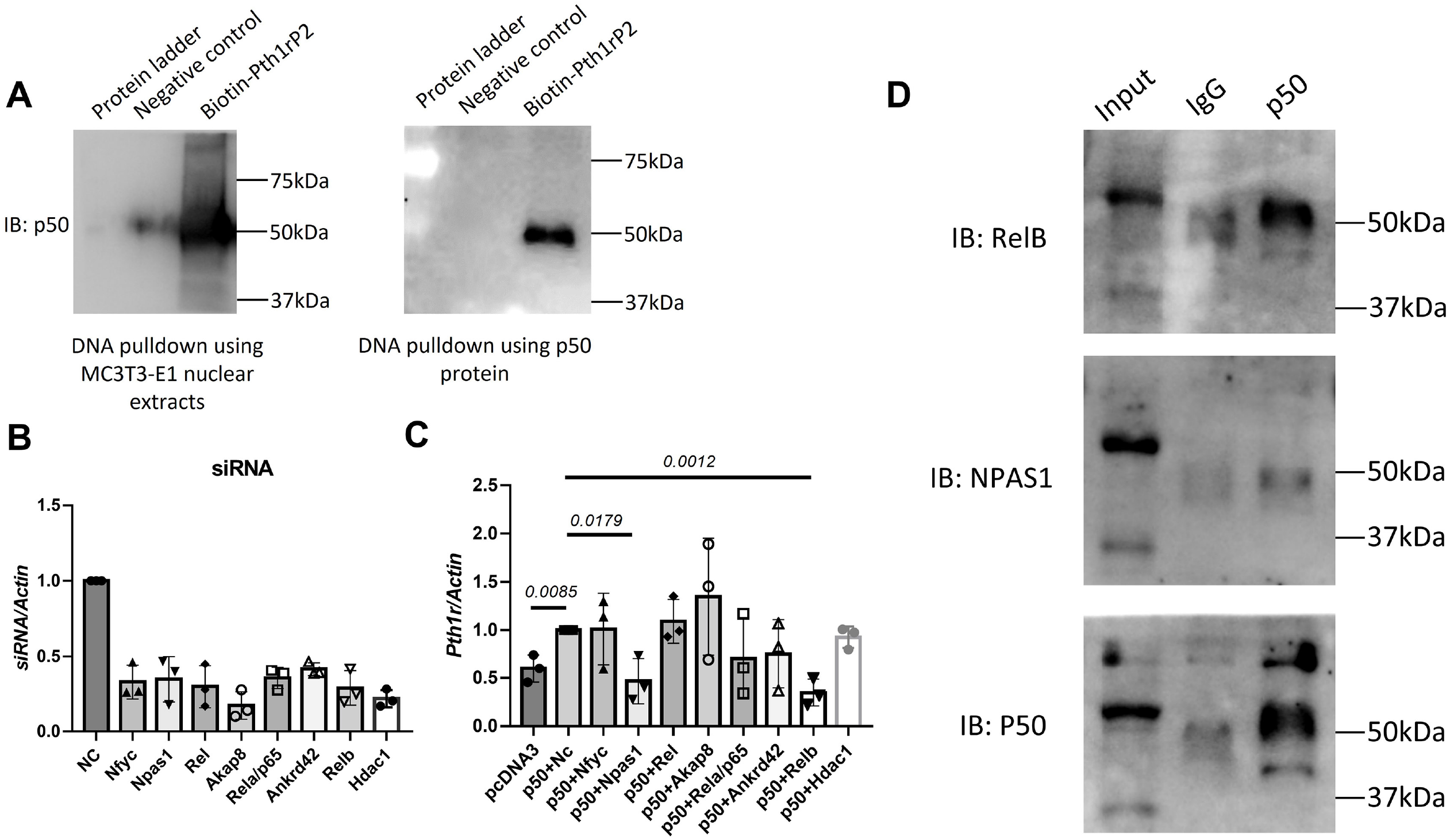
p50 heterodimerize with RelB to transactivate the expression of *Pth1r*. (A) DNA pulldown assay with biotin-labeled Pth1r P2. MC3T3-E1 nuclear extracts or p50 recombinant protein was probed with biotin-Pth1rP2 and then subjected to Immunol blotting using p50 antibody. (B) qPCR results of the expression levels of *Nfyc, Npas1, Rel, Akap8, Rela, Ankrd42, Relb* and *Hdac1* in related siRNA treated MC3T3-E1 cells. Data shown as mean ± SD by one-way ANOVA, n=3 independent experiments for each group. (C) qPCR results of the expression levels of *Pth1r* in p50 overexpression plasmid and *Nfyc, Npas1, Rel, Akap8, Rela, Ankrd42, Relb*, or *Hdac1* siRNA co-transfected MC3T3-E1 cells. (D) IP results using p50 antibody in MC3T3-E1 protein extracts, IgG was used as a negative control.

### *Zfp467 -/-* cells have increased PTH signaling and higher extracellular acidification rates (ECAR)

As the PKA pathway is considered to be one of the major downstream pathways of the PTH signaling network, it would be important to know whether *-/-* cells have higher PKA activation levels due to the higher expression of PTH1R. PTH increased cAMP within 10 minutes, measured by ELISA in +/+ COBs, and *-/-* cells had a higher levels of cAMP expression than +/+ cells (*p*=0.0007 for vehicle, *p*<0.0001 for 50nM, *p*=0.0008 for 100nM). In addition, *-/-* BMSCs also had significantly higher levels of intracellular cAMP after 10-60 min exposure of PTH (*p*<0.0001 for 10 min-100nM, *p*=0.0341 for 30 min-100nM, *p*<0.0001 for 60 min-100nM) (**Fig. 7A, B**). Importantly, -/- COBs and BMSCs showed a greater magnitude of increase of cAMP after PTH treatment (*p=0*.*0053* for interaction in COBs, *p=0*.*0010* for interaction in BMSCs), which resulted from higher PTH1R in -/- cells. Additionally, as CREB was one of the major downstream targets of PKA, we found higher phosphate ratio of CREB in -/- COBs and BMSCs whereas the total protein level of CREB showed no difference (**Fig. 7C, D**).

**Fig. 7.**
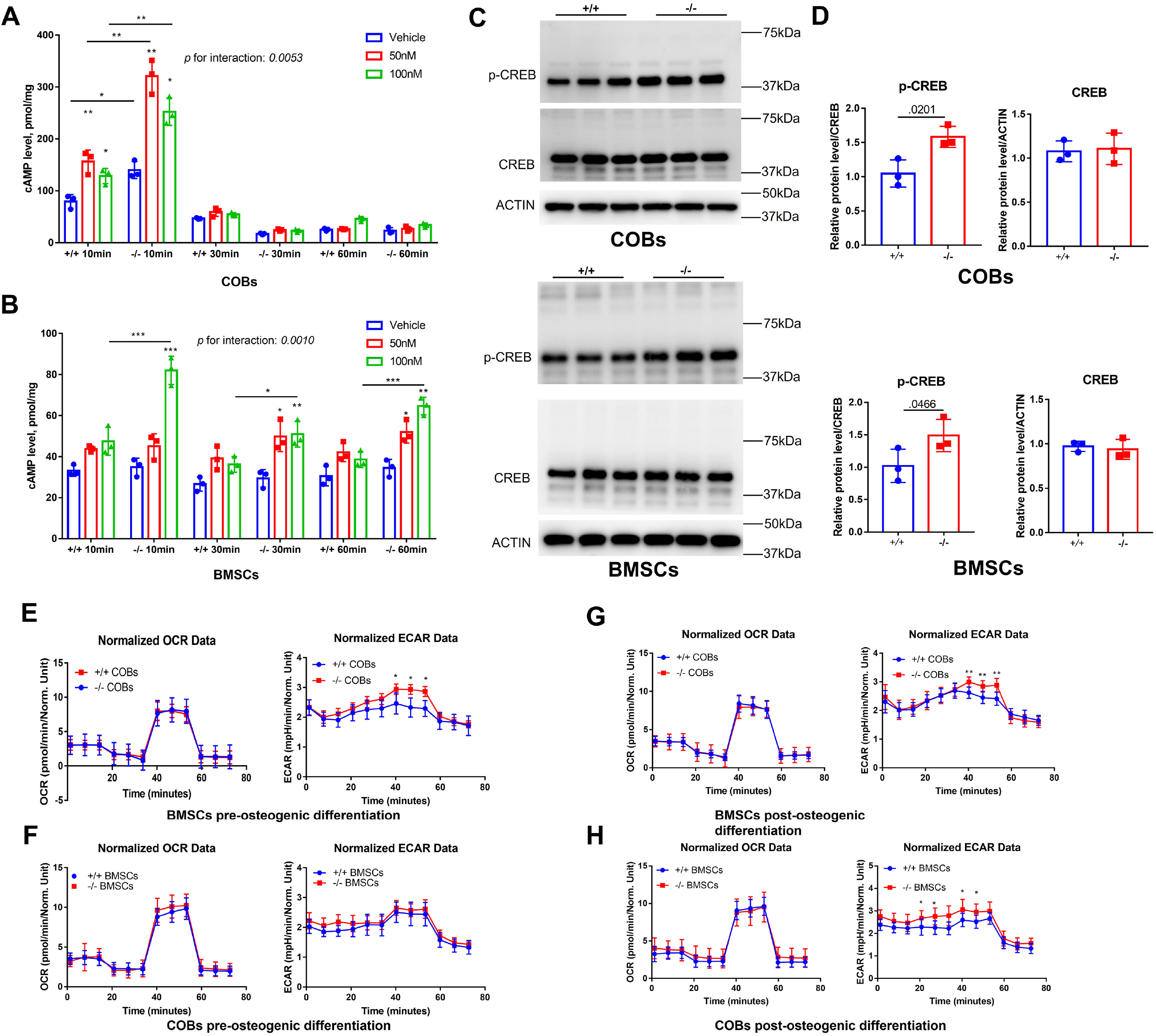
*Zfp467 -/-* cells have increased PTH signaling and higher extracellular acidification rates. (A) Cyclin adenosine monophosphate (cAMP) ELISA of undifferentiated COBs, from 10-60 min with 100 nM PTH treatment, PTH increased cAMP expression within 10 minutes of treatment, and *-/-* cells have higher level of cAMP than *+/+*. Data shown as mean ± SD by two-way ANOVA, n=3 independent experiments for each group; *, *p<0*.*01*, **, *p<0*.*001*. (B) cAMP ELISA of undifferentiated BMSCs, from 10-60 min with 100 nM PTH treatment, *-/-* BMSCs had significantly higher level of intracellular cAMP after 10-60 min exposure of PTH. Data shown as mean ± SD by one-way ANOVA, n=3 independent experiments for each group; *, *p<0*.*01*, **, *p<0*.*001*. (C, D) Western blot and quantitative analysis of pre-differentiated COBs and BMSCs. Higher expression levels of p-CREB but not total CREB was found in both *-/-* COBs and BMSC. Data shown as mean ± SD by unpaired Student’s t test, n=3 independent experiments for each group. (E, F) Oxygen consumption rates (OCR) and Extracellular Acidification Rates (ECAR) of undifferentiated *+/+* and *-/-* COBs or BMSCs. No difference was found regarding OCR between genotypes, but *-/-* BMSCs had higher ECAR than *+/+* BMSCs. Data shown as mean ± SD by unpaired Student’s t test, n=12 technical replicates. *, *p<0*.*01*, **, *p<0*.*001*. (G, H) Oxygen consumption rates (OCR) and Extracellular Acidification Rates (ECAR) of *+/+* and *-/-* COBs or BMSCs after 3 days’ osteogenic differentiation. No difference was found regarding OCR between genotypes, but both -/- COBs and BMSCs cells had significantly higher ECAR level than *+/+* cells. Data shown as mean ± SD by unpaired Student’s t test, n=12 technical replicates. *, *p<0*.*01*, **, *p<0*.*001*.

In a previous study, PTH was shown to enhance aerobic glycolysis, which is a major source of ATP for osteoblast differentiation(16). We measured the oxygen consumption and extracellular acidification in both COBs and BMSCs pre-osteogenic differentiation and 3 days after osteogenic differentiation. Cellular respiration measurements showed that Zfp467-/-pre-differentiated BMSCs had significantly increased ECAR, but no difference was found for pre-differentiated COBs in respect to either ECAR or oxygen consumption rate (OCR) (**Fig. 7E, F**). However, both - /- COBs and BMSCs cells had significantly higher ECAR levels after 3 days of osteogenic differentiation, although no difference was found regarding OCR between genotypes (**Fig. 7G, H**). These data suggest that -/- COBs or BMSCs may have higher glycolysis as a result of higher PTH1R expression.

### *Zfp467-/-* cells showed increased sensitivity to PTH and enhanced pro-osteogenic as well as anti-adipogenic effects

To determine the sensitivity of Zfp467-/-cells to PTH, we treated COBs with osteogenic differentiation media and PTH for 7 or 14 days simultaneously. A dose response to PTH led to a significant increase in ALP staining in *-/-* COBs, and in +/+ cells. Furthermore, 100nM PTH in *-/-* cells produced remarkably higher positive-stained cells than vehicle group (*p*=0.0001) while there was no statistical significance seen among *+/+* groups (*p*=0.6536) (**Fig. 8A, B**). Importantly, -/- COBs showed a more significant response to PTH regarding ALP staining (*p*=0.0201) (**Fig. 8B**). Alizarin red staining showed a parallel trend as ALP staining; an increase in PTH dose resulted in an increase in mineralization for *-/-* COBs only (**Fig. 8A**).

**Fig. 8.**
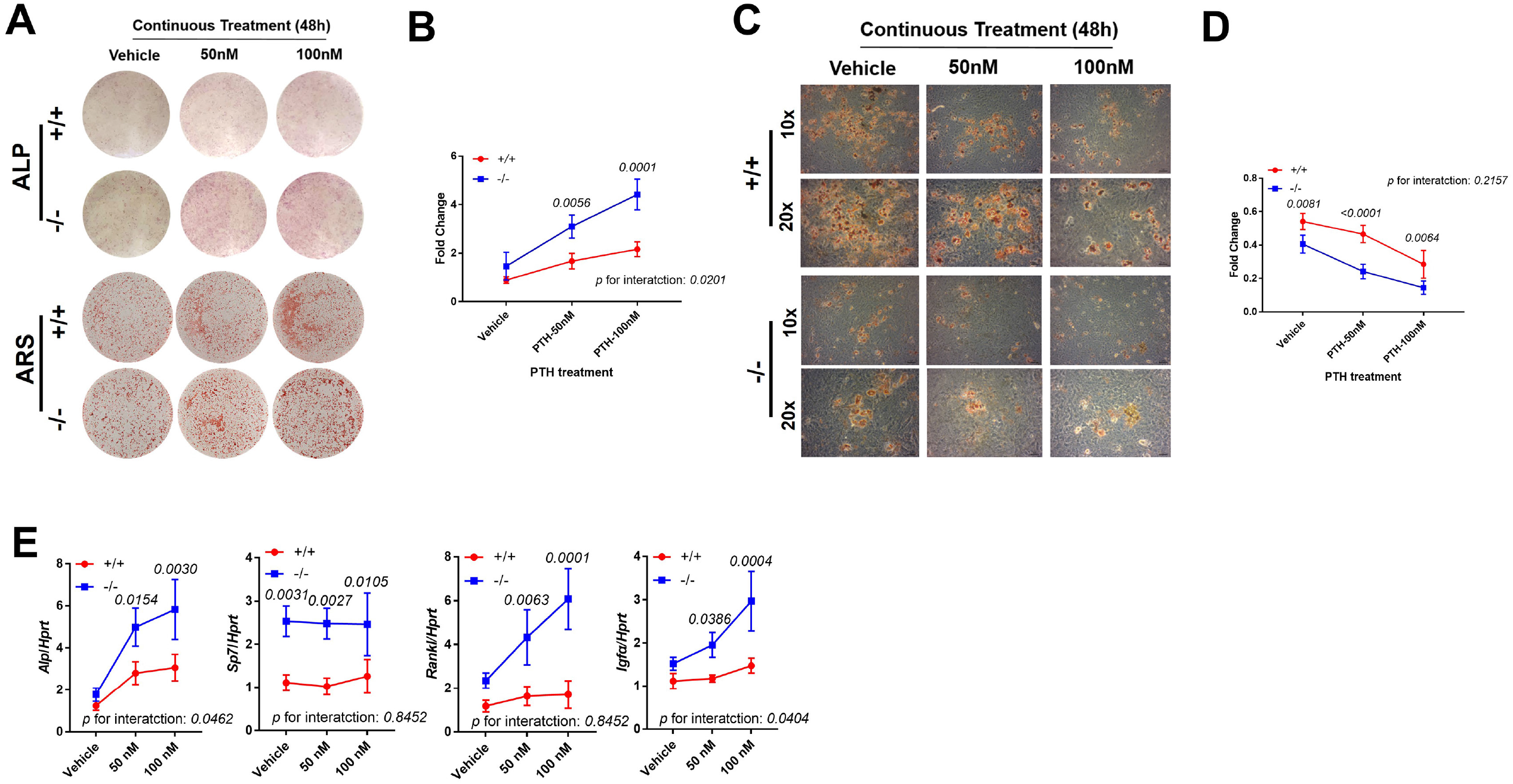
*Zfp467-/-* cells showed increased sensitivity to PTH and enhanced pro-osteogenic as well as anti-adipogenic effects. (A) Representative images of ALP (day 7) and Alizarin red staining (day 14) of differentiated COBs with PTH treatment. An increase in PTH-dose led to an increase in ALP staining and mineralization in both *+/+* and *-/-* COBs. *-/-* COBs showed more ALP positive cells and mineralization than *+/+* COBs. (B) ALP stain quantification in COBs. Data shown as mean ± SD by two-way ANOVA, n=3 independent experiments for each group. (C) Representative images of Oil Red O staining (10X and 20X) of COBs after 14 days in’ osteogenic differentiation with PTH treatment. While PTH treatment inhibited adipocyte formation in *+/+* and *-/-* groups, the *-/-* group showed fewer adipocytes in all treatment groups as compared to *+/+*. (D) ORO stain quantification in COBs. Data shown as mean ± SD by two-way ANOVA, n=3 independent experiments for each group. (E) qPCR results for osteoblast related genes after 7 days’ osteogenic differentiation in COBs. Data shown as mean ± SD by two-way ANOVA, n=3 independent experiments for each group.

In osteogenic media, calvarial progenitors can differentiate into adipocytes. We noted significantly less lipid droplets in -/- adipocytes cultured in osteogenic media compared to untreated *+/+* samples (**Fig. 8C**). Additionally, a reduction in adipogenesis was observed with PTH exposure at 50nM and 100nM in both *+/+* and *-/-* cells, and -/- cells showed much less lipid droplet formation compared to +/+ group (**Fig. 8C, D**). However, the magnitude of decrease after PTH treatment is almost identical in +/+ and -/- cells (**Fig. 8D**). These results were then confirmed by qRT-PCR after 7 days osteogenic differentiation which indicated higher expression of osteogenic differentiation related genes such as *Alp, Sp7, Rankl and Igf-1* in *-/-* compared to *+/+* COBs with PTH exposure (**Fig. 8E, F**). It is noteworthy that there is a statistically significant interaction between genotype and PTH treatment, suggesting an increased sensitivity to PTH in -/- cells.

### Gene Silencing of *Pth1r* or PKA inhibitors suppressed *Zfp467*-/- induced increased osteogenic differentiation

To determine whether the increase in osteogenic differentiation seen in *Zfp467-/-* cells is due to higher PTH1r levels, we knocked down *Pth1r by siRNA* and found that *Pth1r* knock down led to decreased osteogenic differentiation in both +/+ and -/- COBs. Although -/- COBs still show slightly higher *Alp* staining (**Fig. 9A**), *Pth1r* knock down in -/- cells dampens the increase in *Alp* and *Sp7* gene expression during osteogenic differentiation compared to +/+ cells, indicating that this increase is associated with the up-regulation of *Pth1r* seen in -/- cells (**Fig. 9B**).

**Fig. 9.**
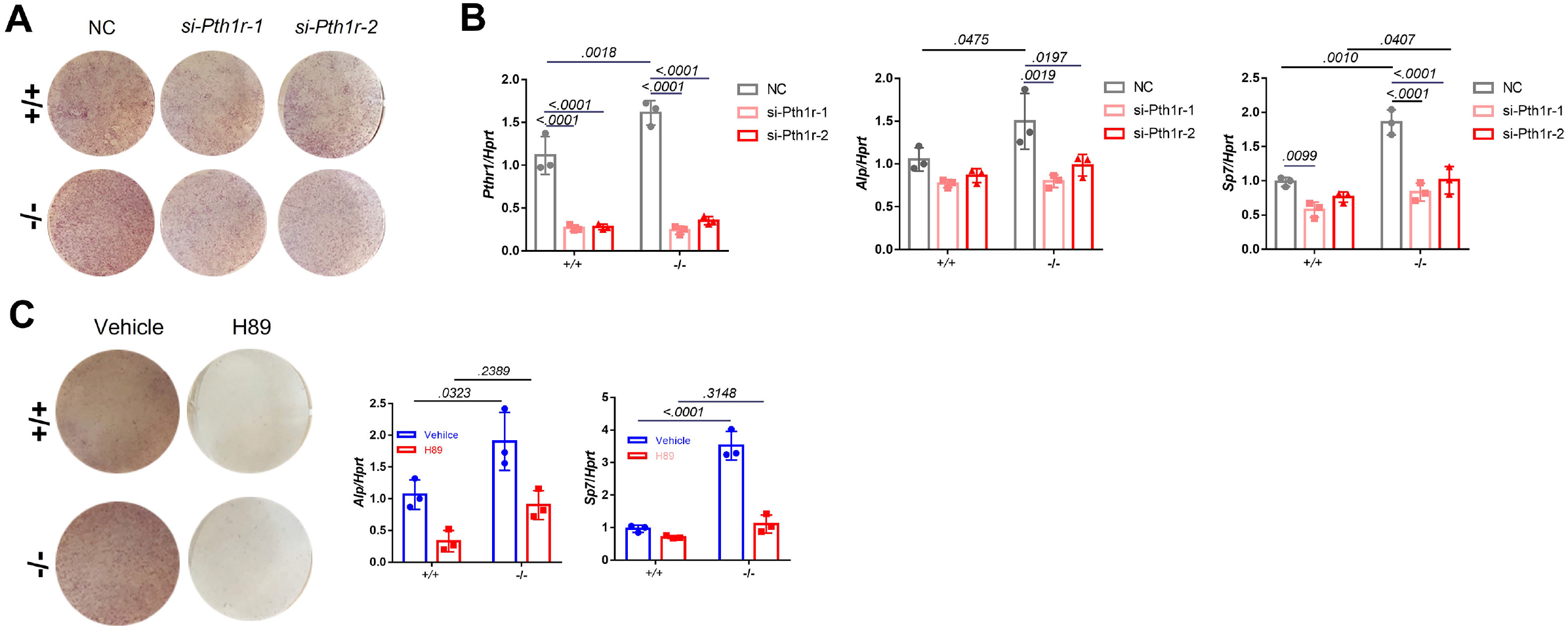
Gene Silencing of *Pth1r* or PKA inhibitors suppressed *Zfp467*-/- induced increased osteogenic differentiation. (A) Representative images of ALP staining of differentiated COBs with *Pth1r* or control siRNA treatment. (B) qPCR results for osteogenic differentiation related genes after 7 days’ osteogenic differentiation and siRNA treatment in COBs. Data shown as mean ± SD by two-way ANOVA, n=3 independent experiments for each group. (C) Representative images of ALP staining and qPCR results for osteogenic differentiation related genes after 7 days’ osteogenic differentiation and PKA inhibitor treatment in COBs. Data shown as mean ± SD by two-way ANOVA, n=3 independent experiments for each group.

Furthermore, a PKA inhibitor could also significantly decrease osteogenic differentiation in COBs (**Fig. 9C**). The staining results showed ALP activity was much higher in -/- cells in the absence of the PKA inhibitor, but no difference was found with the PKA inhibitor between genotypes (**Fig. 9C**). In addition, we observed that with the PKA inhibitor, upregulated osteogenic genes in -/- cells including *Alp* and *Sp7* could be totally reversed (**Fig. 9C**). Similarly, treating COBs with the PKA inhibitor during PTH treatment simultaneously for 7 days led to a suppression of osteogenic differentiation in both +/+ and -/- cells (**Fig. 9-figure supplement**). qPCR results showed that PKA inhibitor could totally reverse the upregulation of *Alp, Sp7* and *Rankl* in PTH treated *-/-* cells (**Fig. 9-figure supplement A, B**).

### *Wdfy1, So×10*, and *Ngfr* and related downstream targets of ZFP467

We then next asked what were the downstream targets of Zfp467. In an unbiased analysis, we used RNAseq in pre- and post-differentiated calvarial osteoblasts from +/+ and *Zfp 467 -/-* cells to assess potential regulatory pathways and differentially expressed genes (**Fig. 10A-D**). The PI 3-K and MAPK signaling pathways were differentially up regulated in Zfp467-/- cells whether pre- or post-differentiated when compared to WT (**Fig. 10D**). There were several highly expressed genes in the -/- cells related to osteogenesis, including *Wdfy1, So×10*, and *Ngfr* (**Fig. 10B**) These results were confirmed by qRT-PCR (**Fig. 10E**). When *Zfp 467* was over-expressed in MC3T3-E1 cells those three genes were significantly suppressed relative to *Gfp* overexpression.

**Fig. 10.**
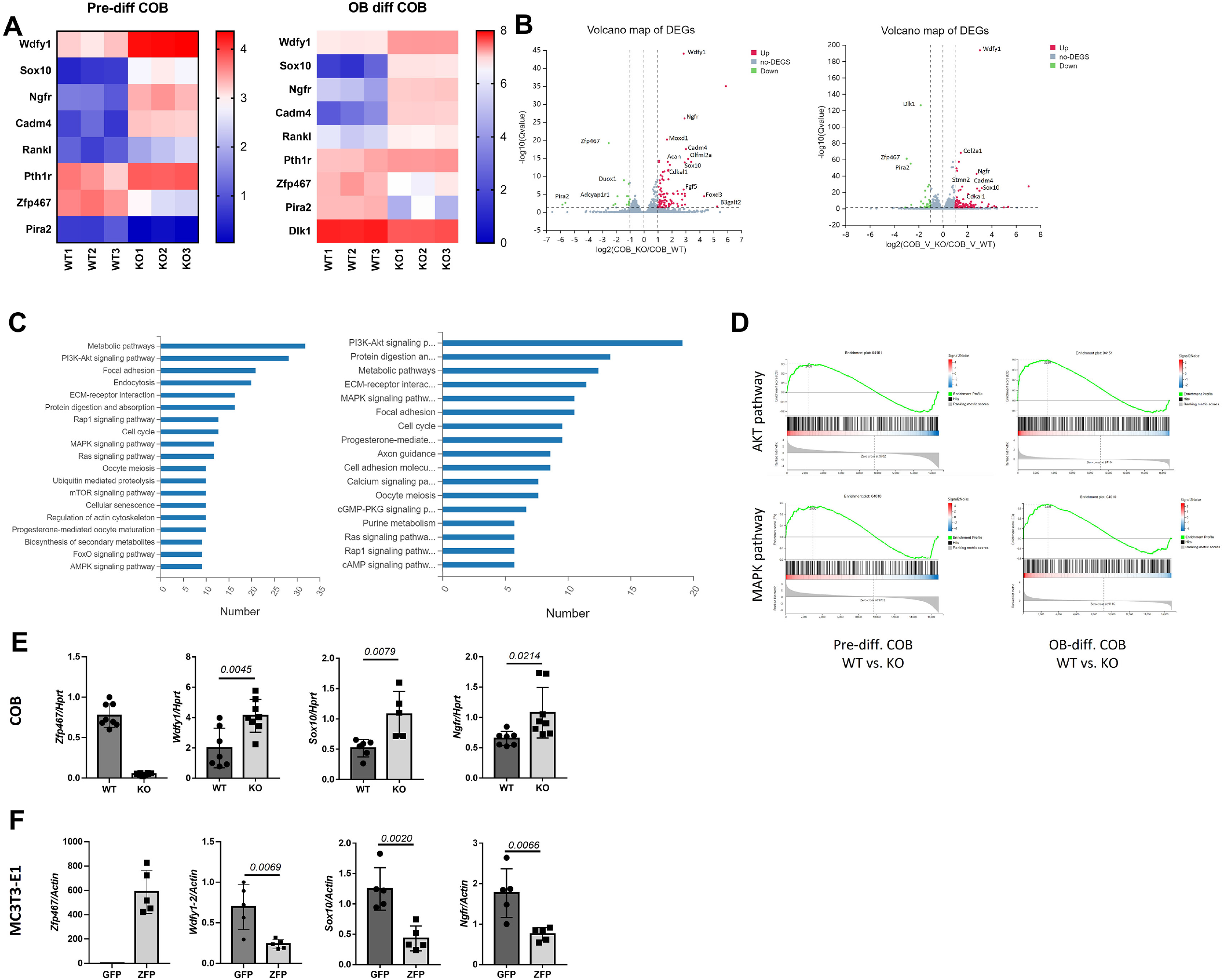
*Wdfy1, So×10*, and *Ngfr* were found upregulated and MAPK, AKT pathways were found activated in *Zfp467 -/-* COBs. (A) Heat map of differentially expressed genes (DEG) from calvarial osteoblasts (+/+ and Zfp467-/-) at pre-differentiation (Panel A) and at differentiation with a p value < 0.05 and a fold change > 2.0 or < -2.0. (B) These DEGs that represented by volcano plots. (C) Functional annotation (Cellular Component [CC]) for the DEGs for +/+ vs -/- for pre- (left) and post-differentiated calvarial OBs (right). (D) GSEA enrichment plots for AKT and MAPK pathways. (E) qRT-PCR was performed on calvarial osteoblasts from +/+ and -/- to confirm gene expression changes noted by RNA seq. (F) Over-expression of *Zfp 467* in MC3T3-E1 cells confirmed statistically significant suppression of the top 3 genes (*Wdfy1, So×10, and Ngfr)*, when compared to gfp over-expression; p<0.05 or lower. Data shown as mean ± SD by unpaired Student’s t test, n=5-8 independent experiments for each group.

## Discussion

In this paper we characterize the role of Zfp467 in the molecular, cellular and biochemical responses to PTH in osteoblast progenitors. Based on our data, we propose a novel pathway by which PTH can enhance its own anabolic actions in osteoblast progenitors by repressing Zfp467, a negative regulator of PTH1R expression. *Zfp467* was originally isolated from mouse hematogenic endothelial LO cells as an OSM-inducible mRNA, which encodes protein ZFP467(17), also called EZI. Zinc finger motifs are involved in protein-protein interactions and protein-DNA binding(10,18,19). Based on the zinc finger domain, zinc finger proteins are classified into C2H2, C3H and C4(20). ZFP467 belongs to C2H2 zinc finger protein family whose zinc finger domain consists of two cysteine and two histidine residues. ZFP467 was initially found to cooperate with STAT3 and augment STAT3 activity by enhancing its nuclear translocation in a kidney cell line(17). ZFP467 is expressed ubiquitously and may be an important mediator of bone marrow cell differentiation into the adipogenic or osteogenic lineages as well as functioning in other tissues in distinct ways (12,13,21).

In our previous work we found that deletion of *pth1r* in mesenchymal cells resulted in low bone mass, high marrow adiposity, and upregulation of *Zfp467*(8). In contrast, the absence of *Zfp467* resulted in a significant increase in trabecular bone volume fraction, accompanied by a marked reduction in peripheral and marrow adipose tissue and improved glucose tolerance(14). Most importantly for the current study, whole bone marrow gene expression by qRT-PCR revealed a ∼40% increase in *Pth1r* expression and greater protein levels in the *Zfp467* -/- mice compared to controls(14). Therefore, we hypothesized that PTH1R and ZFP467 could be involved in a feedback loop whereby the suppression of *Zfp467* mediated by PTH leads to an increase of PTH1R, and therefore an enhanced response to PTH treatment. However, due to the global nature of the Zfp 467 deletion, we could not exclude a cell non-autonomous effect. In the present study therefore we tested that hypothesis both in vivo and in vitro, and sought to determine the cellular and biochemical mechanisms involved in this novel regulatory pathway.

First, we were able to show that conditional deletion of Zfp 467 by PrrxCre led to an increase in osteogenesis and bone formation, a finding that mirrors our results with the global deletion of Zfp 467. On the other hand, using an AdipoCre driven system, we found no effect of conditional deletion in adipocytes on body composition, fat mass, bone mass or total weight. Taken together these data would suggest that Zfp467 is an early mesenchymal transcriptional factor that regulates lineage allocation in early osteoblast but not adipocyte progenitors. Second, in this report, we found constitutive up-regulation of *Zfp467* when we genetically knocked down *Pth1r* in calvarial osteoblasts and bone marrow stromal cells. In addition, much like the study of Quach *et al*.(12), we noted that acute PTH treatment could significantly suppress gene expression of *Zfp467* in both COBs and BMSCs, and the suppression could be partly rescued by both a PKA pathway inhibitor and a PKC inhibitor (12). Similarly, Forskolin, a PKA pathway activator could also inhibit the expression of *Zfp467*, with a more sustained effect. These data indicated that PTH might suppress *Zfp467* expression via activation of the PTH1R through predominantly PKA pathways.

To test the validity of our hypothesis, we investigated how deletion of *Zfp467* affected the expression of the PTH1R. We first examined which transcript of *Pth1r* was upregulated and further confirmed that via dual-fluorescence reporter assay that both P1 and P2 promoters of *Pth1r* were activated in *Zfp467* null cells; however P2 was more activated than P1. Using three different transcription factor prediction databases including PROMO, JASPAR and Animal TFDB, we found several candidate transcription factors that might be involved in the regulation of *Pth1r* in *Zfp467 -/-* cells via activation of the P2 promoter. After overexpressing or knocking down each candidate transcription factor, we found that only p50 could up-regulate *Pth1r* via activation of the P2-2 Pth1r promoter. Further confocal immuno-florescence and nuclear protein detection indicated that the nuclear translocation of p50 was much higher in *Zfp467*-/-cells. Moreover, ChIP-qPCR results showed that p50 could bind to the P2 promoter of the Pth1r. Taken together, these data suggested that the deletion of *Zfp467* resulted in higher nuclear translocation of p50 which bound to the P2 promoter of Pth1r and promoted its transcription.

p50 is one of the DNA binding subunits of the NF-kappa-B (NFκB) protein complex. NFκB is a transcriptional regulator that is activated by various stimuli including cytokines, bacteria and oxidation. The NFκB pathway is involved in several biological processes including inflammation, bone resorption, aging and cancer(22). Activated NFκB translocates into the nucleus and stimulates the expression of an array of genes. However, we found no previous studies that p50 may regulate the gene expression of *Pth1r* or other osteogenic related genes. Nevertheless, it is clear that p50 is involved in osteoclastogenesis and bone resorption, hence it plays a significant role in bone remodeling. It is also conceivable that p50 could associate with histone deacetylase-1 (HDAC1) or be regulated by a PKA catalytic subunit which is also downstream of PTH signaling (23). Moreover, it is likely that other proteins like RelB that might bind to p50 and enhance its effect on the transcriptional regulation of *Pth1r* play a critical role in PTH mediated osteogenesis. Osteoblasts require substrates for energy utilization during collagen synthesis and mineralization. Previous reports by *Lee et al*. and *Guntur et al*. demonstrated that glycolysis is a major source of ATP for differentiating osteoblasts(9,16). Consistently, we found that *Zfp467 -/-* cells showed higher cAMP levels in response to PTH as well as higher glycolytic activity (i.e. ECAR). COBs from *Zfp 467-/-* mice also showed greater differentiation in osteogenic media but less differentiation into adipocytes compared to controls. Higher rates of osteogenesis, enhanced *Rankl* and *Igf1* expression, increased glycolysis and decreased adipogenesis are likely related to greater activation of the PTH1R, possibly due to its higher endogenous expression level.

Since it was reported that the downstream effects of PTH could be partially blocked by PKA inhibitors(24), we used *Pth1r* siRNA and PKA inhibitors to block the effect of PTH1R. We confirmed that PTH1R and its downstream pathways were involved in the deletion of *Zfp467* induced osteogenic differentiation. ALP staining and qPCR results suggested that *Pth1r* siRNA and PKA inhibitor could nearly reverse the upregulated osteogenic differentiation in *-/-* COBs with or without PTH treatment.

Last but not least, we performed RNA-seq to identify potential downstream targets and pathways of *Zfp467* that might be involved in the regulation of osteogenic differentiation. Among the top 3 upregulated genes in *Zfp467*-/-COBs, *Wdfy1* plays an important role in the innate immune responses by mediating TLR3/4 signaling(25,26), which might be also involved in the regulation of osteoblast metabolism and function(27,28). Another significantly upregulated gene, *Ngfr* in Zfp 467-/-cells was found to be required for skeletal cell migration and osteogenesis, especially during injury repair(29,30). NGFR could also mediate AMP-induced modulation of ERK1/2 and AKT cascades(31), which is consistent with our GSEA pathways enrichment. However, further investigation is required to clarify how ZFP467 regulates *Wdfy1, Ngfr* and their downstream AKT or MAPK pathways.

Despite uncovering a novel regulatory circuit through p50, we recognize there are some limitations to our study. First, it should be noted that we also found that *Zfp467-/-* cells have higher nuclear translocation of GATA1 and overexpression of GATA1 could promote *Pth1r* expression. However overexpression of GATA1 failed to activate the P2-2 promoter of *Pth1r*. Hence, we believe that GATA1 might be another of the regulating transcription factors between ZFP467 and Pth1r, but it likely regulates *Pth1r* expression via binding sites other than those present on P2-2.

Second, we have not determined how Zfp467 binds with p50-RelB heterodimer, and we have not identified the exact genomic sequence required for p50-RelB heterodimer binding at the P2 promoter binding site. Further studies will be needed to address this important mechanistic question. Hence the precise mechanism of action of Zfp467, and the downstream consequences of p50-RelB heterodimer binding on the presumed switch between adipocytes and osteoblasts is still not fully defined.

Last, to our knowledge there must be a break on the activation of a feed forward system, but we have to acknowledge that we have not delineated whereby the fast forward system interacts with β-Arrestin or any other turns off signaling.

In summary, taken together, we demonstrated the importance of the zinc finger protein ZFP467 for lineage allocation in vitro and in vivo, as well as responsiveness to PTH in osteoblast progenitors. Our data also support a novel feed-forward regulatory loop whereby suppression of *Zfp467* mediated by PTH and its downstream PKA pathway leads to an increase of PTH1R via p50, and subsequent enhanced responsiveness to PTH treatment. These findings have significant implications for our understanding of the anabolic effects of PTH on bone.

## Materials and methods

### Reagents

PTH were purchased from Bachem (Torrance, CA). H89, G06983 and Forskolin were purchased from Selleck Chemicals (Houston, TX). *Pth1r, p50* and non-targeting control siRNAs were purchased from Life Technologies (Carlsbad, CA). The expression vector for EBF1, MYOD, cFOS, GATA1, p50 and empty vector were obtained from Addgene: p50 pcDNA3 was a gift from Stephen Smale (Addgene plasmid #20018), pcDNA3-GATA1 was a gift from Licio Collavin & Giannino Del Sal (Addgene plasmid #85693), pCAG-EBF1 was a gift from Elena Cattaneo (Addgene plasmid #96965), pCAG-MyoD was a gift from Andrew Lassar (Addgene plasmid #8398), MMTV-cFOS-SV40 was a gift from Philip Leder (Addgene plasmid #19259). pCMV-Creb1 was a gift from Georgios Stathopoulos (Addgene plasmid # 154942)

### Animals

*Zfp467*^*fl/fl*^ on the C57BL/6 J background was generated by Cyagen (Fig. 1 supplement 1). Exons 2∼4 were selected as conditional knockout region (Transcript: Zfp467-001 ENSMUST00000114561). The targeting vector, homology arms and cKO region were generated by PCR using Bacterial Artificial Chromosome (BAC) clone RP24-144J8 and RP23-24K23 from the C57BL/6J library as template. To generate mice lacking Zfp467 in limb mesenchymal stem cells, *Prrx1Cre; Zfp467*^*fl/fl*^ were generated by crossing *Prrx1Cre* transgenic mice to *Zfp467*^*fl/fl*^ mice (*Prx*-Cre *Zfp467*^*fl/fl*^) and *·*^*fl/fl*^ mice were used as controls. To generate mice lacking Zfp467 in adipocyte tissues, *AdipoCre; Zfp467*^*fl/fl*^ were generated by crossing *Adiponectin Cre* transgenic mice to *Zfp467*^*fl/fl*^ mice and *Zfp467* ^*fl/fl*^ mice were used as controls. Total DNA was isolated from ear punch biopsies, and routine PCR was used to genotype mice.

All experiments were performed with 12 weeks old - and sex-matched littermates. All animals are in the C57/Bl6 background and were housed in polycarbonate cages on sterilized paper bedding and maintained under 14:10 hour light:dark cycles in the barrier, AAALAC-accredited animal facility at Maine Medical Center Research Institute or Harvard Center for Comparative Medicine. All experimental procedures were approved by the Institutional Animal Care and Use Committee of Maine Medical Center and followed the NIH guidelines for the Care and Use of Laboratory Animals and also approved by the Harvard University Institutional Animal Care and Use Committee.

No statistical methods were used to predetermine sample size. Mice label and measurements were performed by two independent researchers and selected at random (by cage) into following experiments. Masking was used during data collection and data analysis. Animals with ulcerative dermatitis or other diseases were excluded from the study.

### Dual-energy X-ray Absorptiometry (DXA)

Whole body composition evaluation of the head was performed using the PIXImus densitometer (GE-Lunar, Fairfield, CT, USA). The PIXImus was calibrated daily with a phantom provided by the manufacturer.

### Micro-computed Tomography (μCT)

A high-resolution desktop micro-tomographic system (vivaCT 40, Scanco Medical AG, Brüttisellen, Switzerland) was used to assess the trabecular and cortical bone microarchitecture, volume and mineral density in mouse femurs. Scans were acquired using a 10.5 μm^3^ isotropic voxel size, 70 kVp peak x-ray tube intensity, 114 mA xray tube current, 250 ms integration time, and were subjected to Gaussian filtration and segmentation. All scans were analyzed using manufacturer software (Scanco, version 4.05). The trabecular bone region of interest started 210 μm (20 transverse slices) proximal to the break in the growth plate and extended 1575 μm (150 transverse slices). Bone was segmented from soft tissue using a mineral density threshold of 375 mg HA/cm^3^. Trabecular bone was analyzed for bone volume fraction (Tb.BV/TV, %), trabecular thickness (Tb.Th, mm), trabecular number (Tb.N, mm-1), trabecular separation (Tb.Sp, mm), connectivity density (Conn.D, mm-3), and trabecular bone mineral density (Tb.BMD, mg HA/cm3). The cortical bone region of interest started at 55% of the total bone length distal to the femoral head and extended 525 μm (50 transverse slices). Bone was segmented using a mineral density threshold of 700 mg HA/cm^3^. Cortical bone was analyzed for bone area (Ct.Ar, mm2), medullary area (Ma.Ar, mm2), bone area fraction (Ct.Ar/Tt.Ar, %), cortical thickness (Ct.Th, μm), cortical tissue mineral density (Ct.TMD, mg HA/cm3), polar moment of invertia (J, mm4) and maximum and minimum moment of inertia (Imax, Imin, mm4).

### Marrow adipose tissue quantification by osmium tetroxide staining and μCT

At the time of sacrifice, tibiae were isolated and placed into 10% neutral buffered formalin overnight at 4°C. Soft tissue was carefully removed to ensure that the fibula remained intact and the bones were washed under continuous cold PBS for one hour, then stored in PBS at 4°C. Quantification and visualization of marrow adipose tissue was performed as described previously(32). Briefly, bones were decalcified in 14% EDTA (pH 7.4) for 14 days, with EDTA changes every 3-4 days. Bones were then washed for 10 minutes in PBS (3 times) and stained with a 1:1 mixture of 2% aqueous osmium tetroxide (cat# 23310-10, Polysciences, Inc., Warrington, PA, USA) and 5% potassium dichromate for 48 hours. Stained bones were then washed with PBS (pH 7.4) for 5 hours (3 times), and subsequently scanned by μCT. Bone marrow adipose tissue (BMAT) content was calculated by determining the whole volume of second osseous center of tibiae.

### Primary cells isolation

The generation of *Zfp467-/-* and wild-type mice was previously described. Calvaria osteoblasts (COBs) were isolated from calvarias of 3-5-day old *+/+* and *-/-* neonates as described in the following protocols(33,34). Bone marrow stromal cells (BMSCs) were isolated from tibiae and femurs of 6-week-old *+/+* and *-/-* female mice as described in the previously established protocols(14,35). All studies were reviewed and approved by the Institutional Animal Care and Use Committee of Maine Medical Center and followed the NIH guidelines for the Care and Use of Laboratory Animals.

### Parathyroid Hormone (PTH) Treatment

Parathyroid hormone 1-34 bovine (PTH, Bachem, Torrance, CA) was made to 1mg/mL stock using a vehicle solution (4mM HCl contained 0.1% BSA). PTH was administered by adding to osteogenic media every 48 hours for each group..

### RNA interference and plasmid transfection

24 hours after seeding, RNA oligos were transfected into COBs or BMSCs using Lipofectamine™ RNAiMAX Transfection Reagent according to the manufacturer’s instructions for 48 hours and the final concentration of siRNA was 5 nM. ORF clone expression vector or a controlled vector was introduced into COBs, BMSCs or MC3T3-E1 cells using Lipofectamine™ 3000 Transfection Reagent for 72 hours once cells reached 80% confluence. All transfection reagents were purchased from Thermo Fisher Scientific, Waltham, MA.

### Real-time PCR and western blot

Total RNA was isolated using a standard TRIzol extraction (Life Technologies, Carlsbad, CA) method. cDNA was generated using the High Capacity cDNA Reverse Transcription kit (Life Technologies, Carlsbad, CA) according to the manufacturer’s instructions for real-time PCR. Proteins from cell culture were extracted by scraping the culture wells in the presence of RIPA buffer (BioRad, Hercules, CA) with protease inhibitor and phosphatase inhibitor (St. Louis, MO). Cytoplasm and nuclear protein were extracted using Nuclear and Cytoplasmic Extraction Reagent Kit (Thermo Scientific, MA, US). Antibodies used for western blot and Chromatin Immunoprecipitation (ChIP) or co-Immunoprecipitation (co-IP) were listed in Table 1.

**Table 1.** Antibody list for co-IP, ChIP and western blot.

| Antibody | Supplier | Cat Num |
| --- | --- | --- |
| ACTIN | Santa Cruz Bio | SC47778 |
| PTH1R | Sigma | SAB4502493 |
| GATA1 | Cell Signaling Technology | 3535T |
| p50 (IP) | Cell Signaling Technology | 13586S |
| p50 (IB) | Cell Signaling Technology | 13681S |
| Histone H3 | Cell Signaling Technology | 9715S |
| RelB | Santa Cruz Bio | sc-48366 |
| NPAS1 | Santa Cruz Bio | sc-376083 |
| p-CREB (SER133) | Cell Signaling Technology | 9198S |
| CREB | Cell Signaling Technology | 9197T |

### Nuclear translocation detection

*Zfp467+/+* and *-/-* BMSCs were plated on an 8-chamber slide (Sigma, St. Louis, MO). Protein location of GATA1 and p50 were detected by immune-florescence using Confocal Microscopy (Leica DMI6000). Nuclear protein was isolated from baseline *Zfp467+/+* and *-/-* BMSCs lysate by Pierce™ NE-PER® Nuclear and Cytoplasmic Extraction Reagent Kit (Life Technologies, Carlsbad, CA). Protein level of GATA1 and p50 were measured using nuclear extracts.

### Dual-fluorescence reporter assay

The Pth1r P1 (0.5 kb: 0-500, 1.1 kb: -600-500; 1.5kb: -1109-500, 2.1kb: -1598-500, P1: -581- -1109) and P2 promoters (P2-1: -449-0, P2-2: -212- -826) were cloned through PCR into the pGL4.20 luciferase reporter vector (Promega, Madison, WI) using C57BL/6 genomic DNA as a template. pGL4.75 luciferase reporter was used as a positive control for transfection efficiency normalization. Baseline COBs, BMSCs or MC3T3-E1 cells were transfected with the reporter constructs and incubated for 48 hours. Luciferase activity was measured using GloMax-20/20 (Promega, Madison, WI). The transcriptional activity was expressed as the ratio of firefly:Renilla luciferase activity

### ChIP Assay and Biotin-*Pth1r* pulldown

MC3T3-E1 subclone 4 (CRL-2593) cell line was purchased from ATCC and used for ChIP and DNA-pull down assay. When MC3T3-E1 cells reached confluency, they were then fixed in 1% paraformaldehyde/PBS for 10 min at room temperature. Chromatin shearing and immunoprecipitation were performed using EZ-Magna ChIP A/G Chromatin Immunoprecipitation Kit (Sigma, St. Louis, MO) according to the manufacturer’s instructions. The immunoprecipitated DNA fragments were used as templates for PCR amplification. ChIP-PCR primers sequences were listed in Table 2. PCR products were used for nucleic acid electrophoresis to avoid unspecific amplification.

**Table 2.**
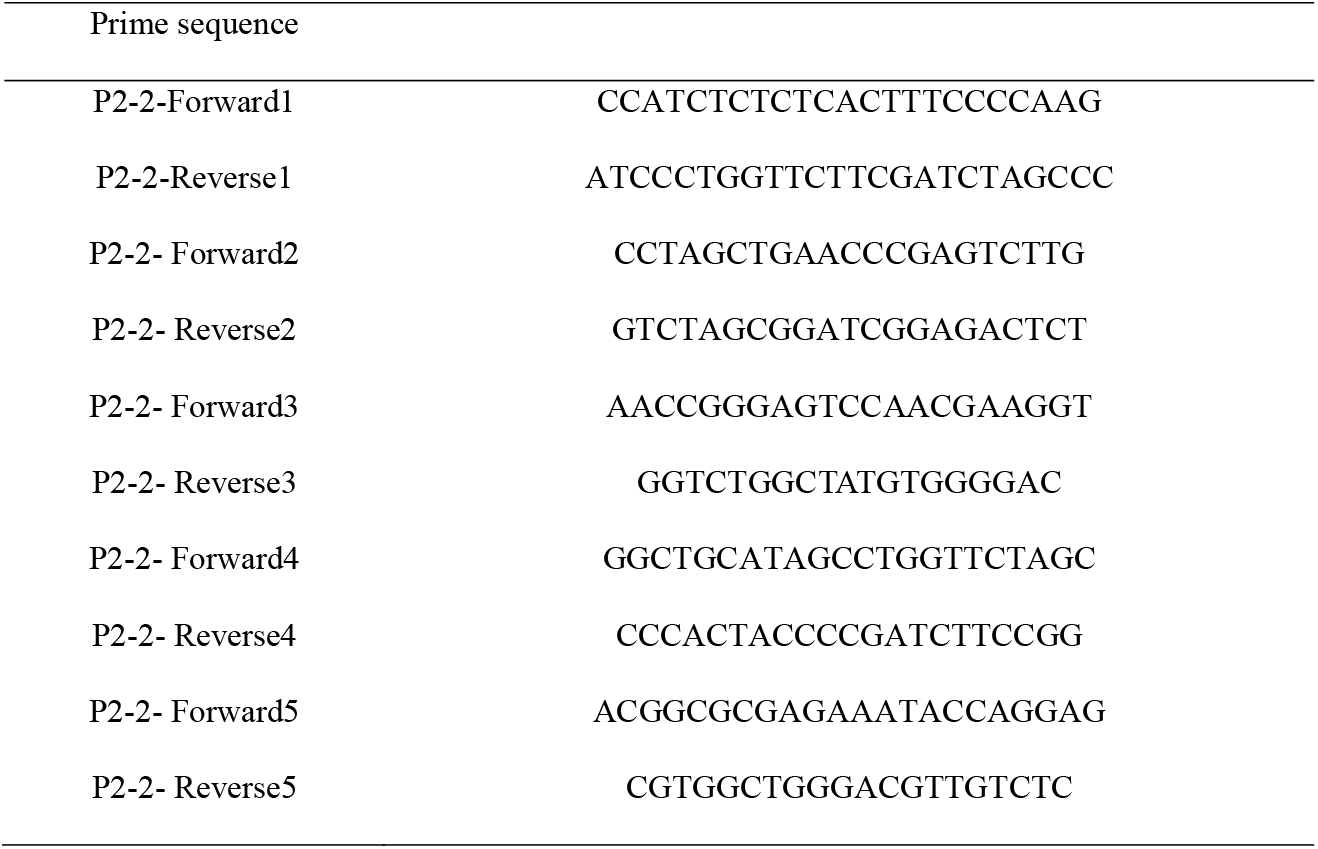
Primer list of ChIP-qPCR for P2-2 promoter of *Pth1r*.

DNA pulldown assay was performed according to previously reported protocol (36). 5’ biotin-modified Pth1r P2-2 dsDNA probe was generated using PCR with oligonucleotide primers modified at its 5’ end by Sangon Biotech (Shanghai, China). Nuclear protein were extracted using Nuclear and Cytoplasmic Extraction Reagent Kit (Thermo Scientific, MA, US). p50 purified protein was purchased from Proteintech (Rosemont, US). Briefly, the Pth1r P2-2 or oligo probe was incubated with streptavidin-coupled Dynabeads (Invitrogen, US) at room temperature for 1 hour to generate probe-bound Dynabeads, and then the probe-bound Dynabeads were incubated with MC3T3-E1 nuclear extracts or p50 purified protein at 4 °C overnight. The protein bound to the probe and beads were eluted and used for gel electrophoresis.

### Determination of Cellular cAMP Levels

Intracellular cAMP expression levels were measured after 10min to 48h of PTH treatment using Mouse/Rat cAMP Assay Parameter™ Kit (R&D Systems, Minneapolis, MN) according to the manufacturer’s instructions in both pre-differentiation COBs and BMSCs. Total protein quantity was used for normalization using the Pierce® BCA Protein Assay Kit (Life Technologies, Carlsbad, CA).

### Cellular Respiration Measurements

The oxygen consumption rate (OCR) and extracellular acidification rate (ECAR) were assessed pre-differentiation or 3 days after osteogenic differentiation using the XF96 Extracellular Flux Analyzer (Seahorse Biosciences, North Billerica, MA), as previously described.(35). Briefly, BMSCs and COBs were plated in 96-well Seahorse XF96 Cell Culture Microplates (Seahorse Biosciences, North Billerica, MA). Several mitochondrial electron transport chain complex inhibitors were used during the test: 2.52uM oligomycin, 12.65 uM carbonylcyanide p-(triflueormethoxy) phenylhydrazone and 12.67 uM Rotenone. All inhibitors were purchased from Seahorse Biosciences. Cell counts via fluorescence detection with Hoechst (Life Technologies, Carlsbad, CA) was used for normalization.

### Osteogenic differentiation and related measurements

*In vitro* osteoblast differentiation and measurement were done according to previously published protocols(14). Briefly, BMSCs and COBs were plated at a density of 1×10^6^/well in 6-well plates. When cells reached around 80% confluency, osteogenesis induction began using osteogenic induction media which consisted of complete αMEM (αMEM, 5% fetal bovine serum, and 1% penicillin/streptomycin), 50ug/mL ascorbic acid, and 8mM beta-glycerophosphate (both were purchased from Sigma, St. Louis, MO)]. αMEM and penicillin/streptomycin were purchased from Life Technologies (Carlsbad, CA), FBS was purchased from VWR (Radmor, PA). Medium was changed every other day until cells were ready to stain for alkaline phosphatase (ALP) and alizarin red (ARS) to assess osteoblasts and mineralization, respectively, around d7 and d14 after differentiation. Additionally, RNA was extracted on day 7 after differentiation for real time PCR analysis.

### Alkaline Phosphatase and Alizarin Red Staining

Alkaline phosphatase staining was performed using ALP kit obtained from Sigma (St. Louis, MO) according to the manufacturer’s instructions at d4 and d7 after osteogenic differentiation for BMSCs and COBs, respectively. ARS (Sigma, St. Louis) staining was done using 1% solution at pH 4.2 at d14 after osteogenic differentiation for both BMSCs and COBs. Briefly, after fixation, cells were stained for 30 minutes with ARS solution at room temperature. Cells were then washed a couple of times with water before they were visualized under the microscope (Leica DM IRB, TV camera). Five randomly fields were chosen to capture images using the ZEISS Efficient Navigation, blue edition camera (Bloomfield, CT) per treatment group for quantification.

### Oil Red O (ORO) staining

Differentiated adipocytes were stained using ORO solution at d9 after adipogenic diff. Briefly, cells were fixed with 10% neutral buffer formalin (Sigma, St. Louis). After that, they were washed with 60% isopropanol (Sigma, St. Louis, MO) before stained with ORO solution (Sigma, St. Louis, MO) for 15 minutes at room temperature. Cells were then washed a couple of times with water before they were visualized and pictures were taking using the Zeiss microscope. For quantification, lipid from adipocyte droplets was extracted using isopropanol and the absorbance was measured at 490 nm using a plate reader (MRX Dynex Technologies, Chantilly, VA).

### RNA-seq and Analysis

RNA-seq was performed on samples from COBs isolated from *Zfp467+/+* and *-/-* mice, pre- or after-4 days’ osteogenic differentiation. RNA-seq analysis was performed by BGI Group (UK, China). Samples were de-identified, and the analysis was blinded to assignment. We employed DAVID 6.8 to perform the functional annotation (Cellular Component [CC]) and GSEA enrichment plots for related pathways. Protein coding genes were assessed with p<0.05, fold change > 2.0 or < -2.0. The enriched CC was evaluated using a false discovery rate (FDR, Benjamini–Hochberg method) of 0.1 and DEG number. We have uploaded our original sequencing data to Sequence Read Archive database (PRJNA877934, http://www.ncbi.nlm.nih.gov/bioproject/877934).

### Statistical Analysis

All data are expressed as the mean ± standard deviation (SD) unless otherwise noted. Results were analyzed for statistically differences using Student’s t-test between two groups or 2-way ANOVA followed by Bonferroni’s multiple comparison post hoc test among three or more groups where appropriate. All statistics were performed with Prism GraphPad 7.0 statistical software (GraphPad Software, Inc., La Jolla, CA). Values of p<0.05 were considered statistically different.

## Acknowledgements

This study was supported by NIDDK R01 DK112374-RB and CJR; NIAMS R01AR073774-RB and CJR; NIDDK R24 DK092759-CJR.

## Data availability

Data are available upon reasonable request.

---

**Fig.1-figure supplement 1.**
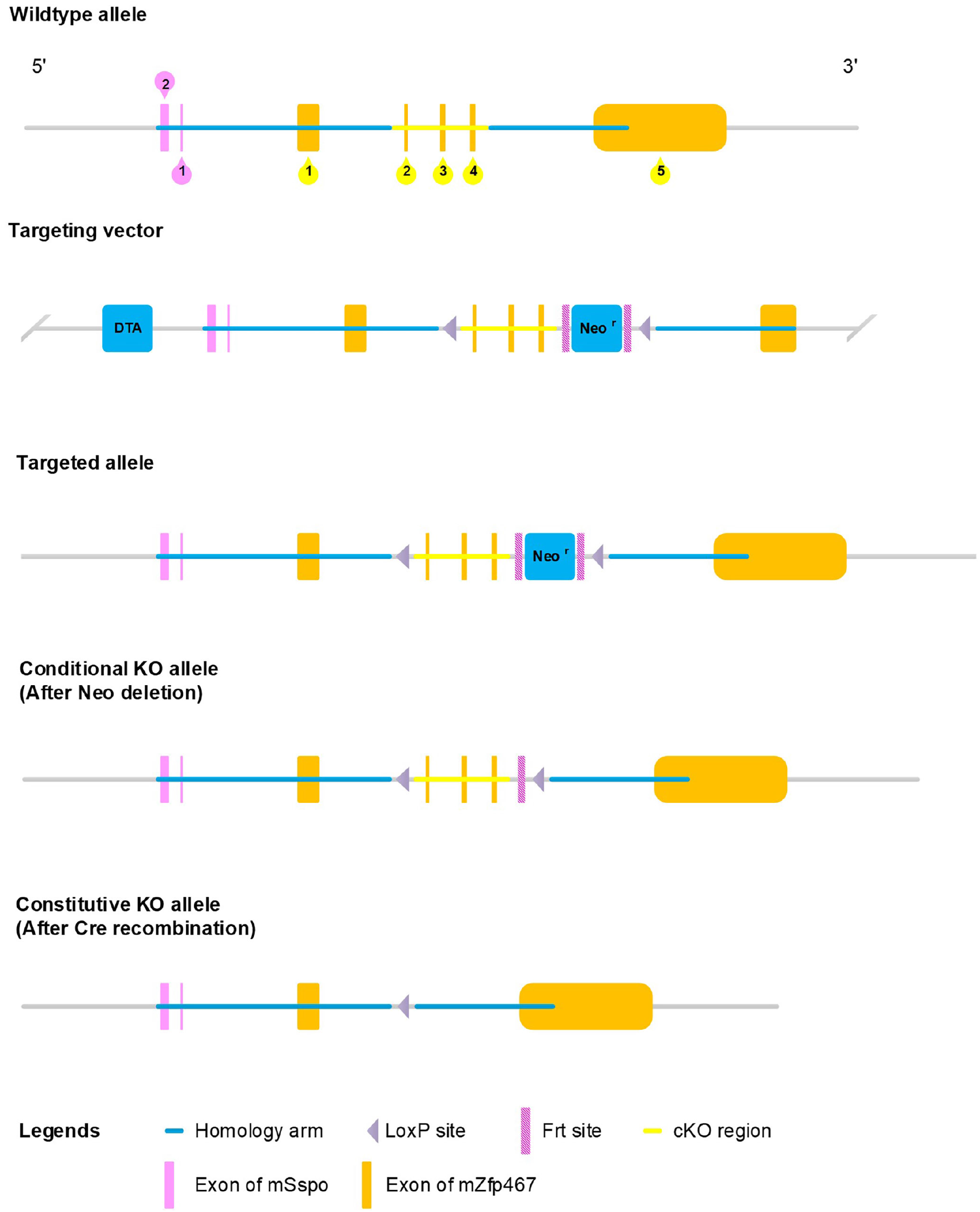
Generation of Zfp467 flox mice. Exons 2∼4 were selected as conditional knockout region. The targeting vector was generated by PCR using BAC clone RP24-144J8 and RP23-24K23 from the C57BL/6J library as template.

**Fig. 1-figure supplement 2.**
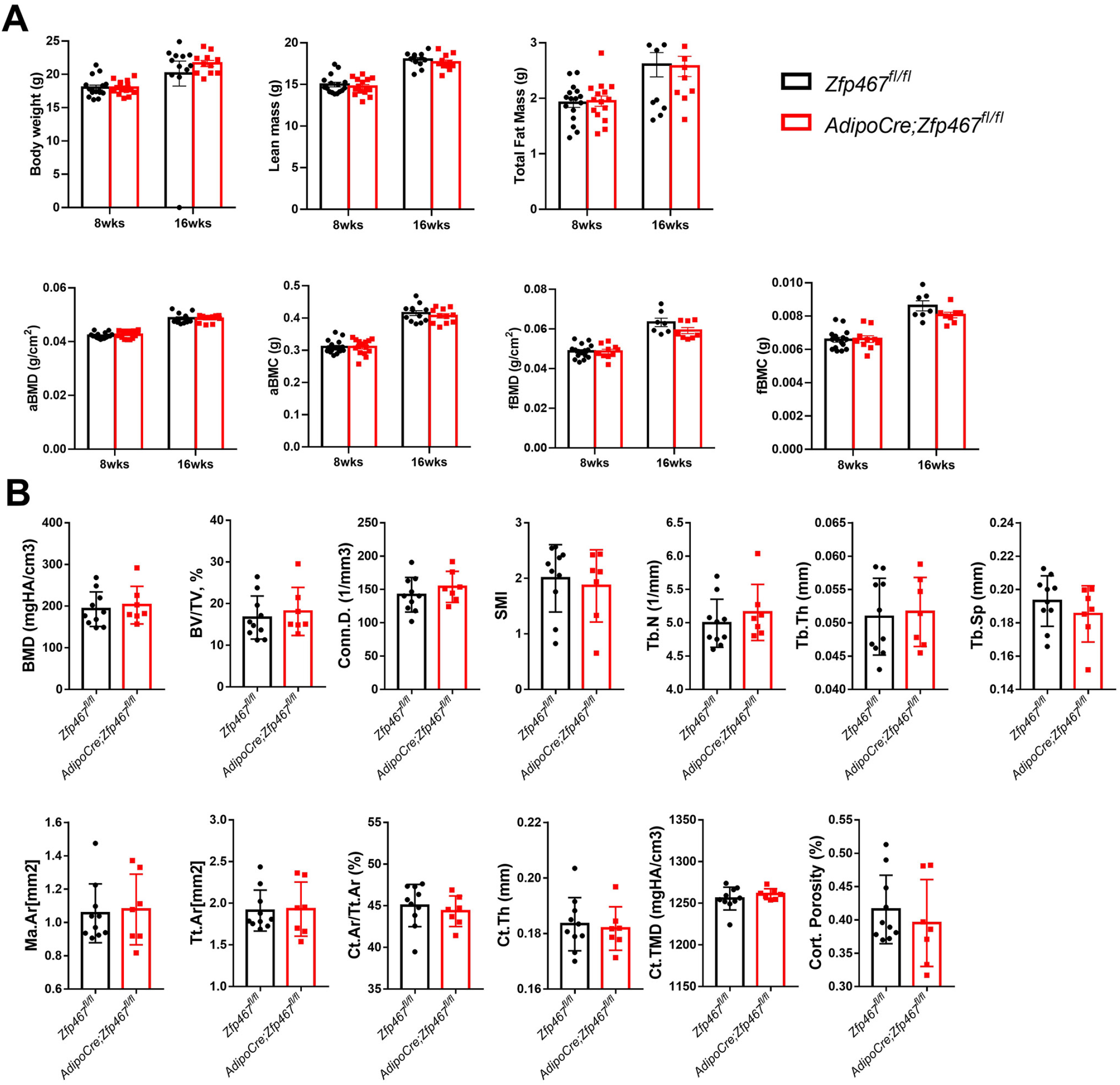
*Adiponectin Cre Zfp467* have similar trabecular and cortical bone mass with controls. (A) Male and (B) Female 12 weeks old mice and control *Zfp467*^*fl/fl*^; *AdipoCre* mice were measured representative μCt images of tibia trabecular and cortical bone of tibiae. Data shown as mean ± SD by unpaired Student’s t test, n=7-10 per group.

**Fig.4-figure supplement.**
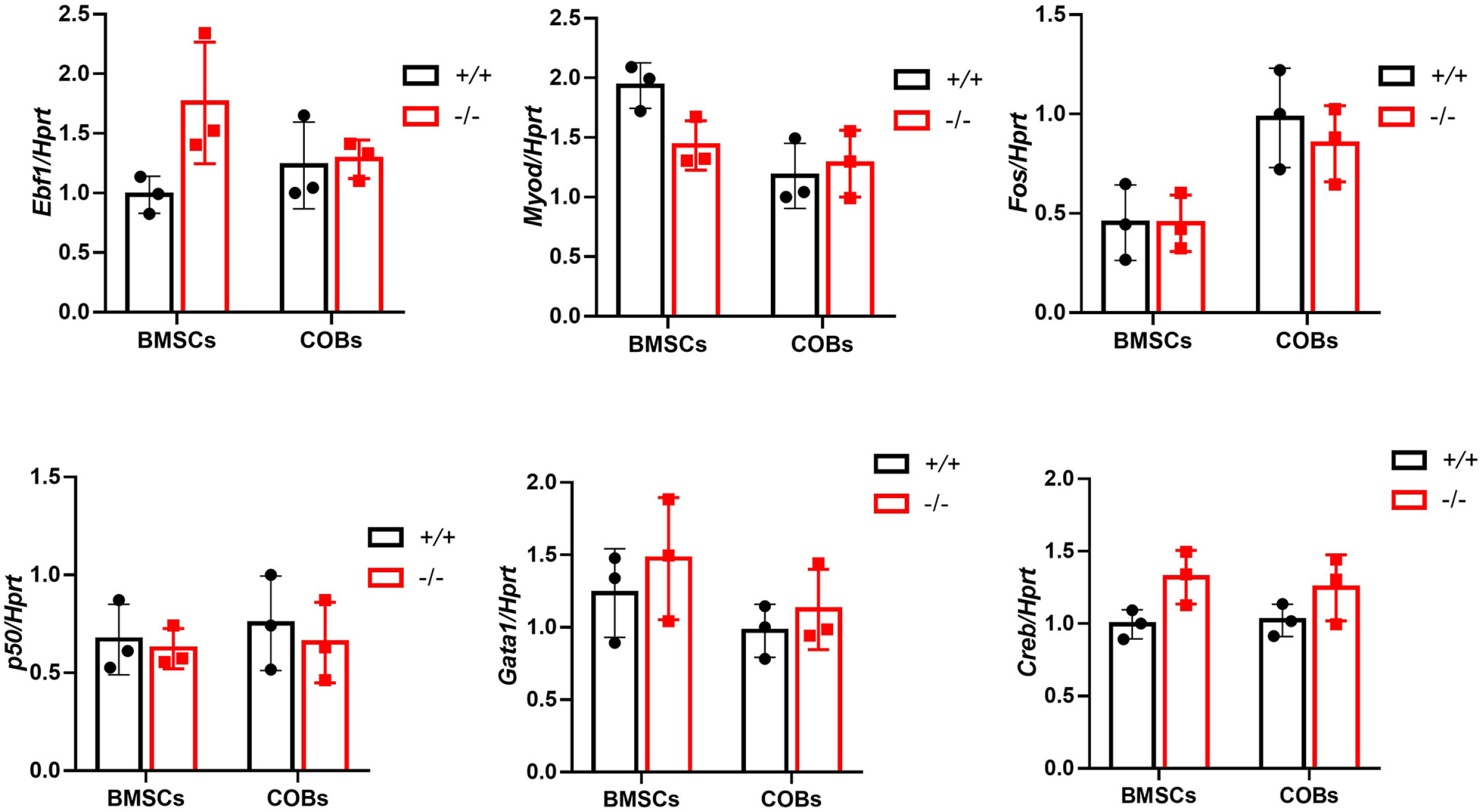
qPCR results of expression of *Ebf1, Myod, Fos, Gata1, p50* and *Creb* in *Zfp467+/+* and *Zfp467-/-* COBs and BMSCs. Data shown as mean ± SD by unpaired Student’s t test, n=3 independent experiments for each group.

**Fig.5-figure supplement.**
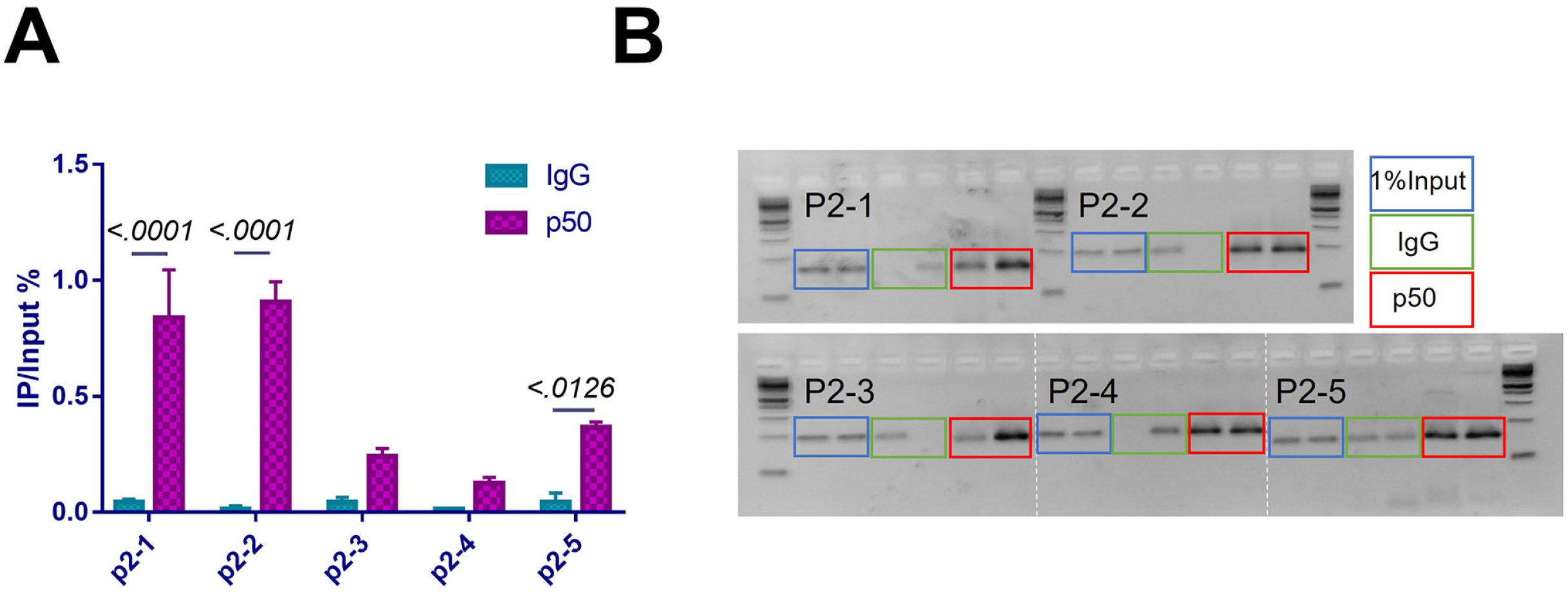
(A) DNA enrichment of Pth1r P2 promoter, ratio between IP and input, first two parts of P2 were enriched using p50 antibody. Data shown as mean ± SD by unpaired Student’s t test, n=3 replicates for each group. (B) RT-PCR product of chromatin immunoprecipitation assay using the nuclear extracts from MC3T3-E1 cells.

**Fig. 9-figure supplement.**
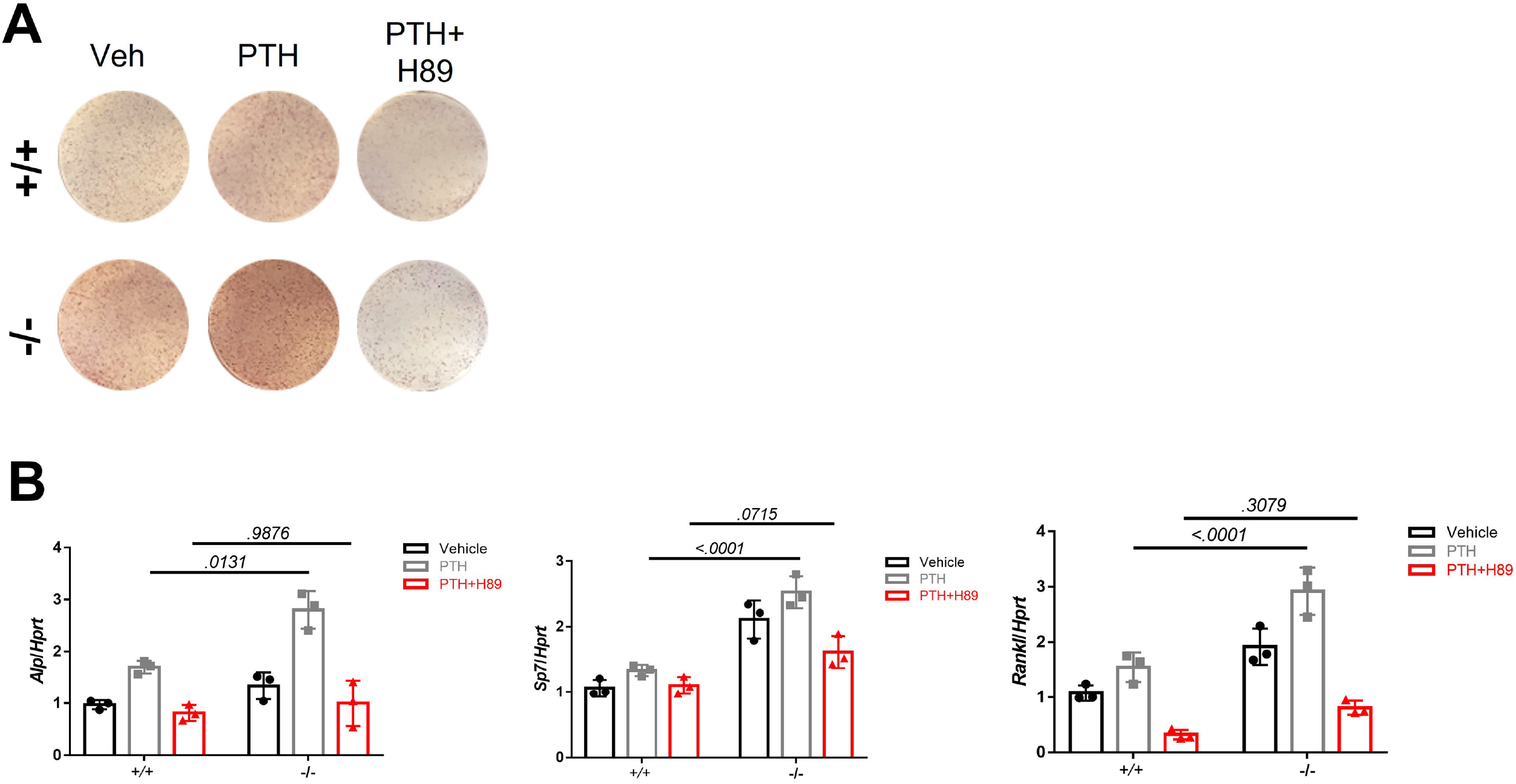
(A) Representative images of ALP staining of differentiated COBs with 100nM PTH treatment and PKA inhibitor. (B) qPCR results for osteogenic differentiation related genes after 7 days’ osteogenic differentiation, PTH treatment and PKA inhibitor treatment in COBs. Data shown as mean ± SD by two-way ANOVA, n=3 independent experiments for each group.

